# AEGIS reveals epitope- and clone-resolved convergence of CNS B and T cell autoreactivity in ROHHAD

**DOI:** 10.64898/2026.05.06.722823

**Authors:** Aaron Bodansky, Vida Ahyong, Monica Dayao, James Asaki, ShiLu Vanasupa, Ravi Dandekar, Shini Chen, Joe Sabatino, Sam Klauer, Giselle Knudsen, Carlos Lizama Valenzuela, Krista McCutcheon, Kelsey Zorn, Sukhman Sidhu, David Yu, Samantha Garcia, Zoe Quandt, Chung-Yu Wang, Bryan Castillo Rojas, Iris Tilton, Y. Rose Citron, Akshay Sharathchandra, Chloe Gerungan, Stuart Tomko, Nathaniel M. Robbins, Andrew McKeon, Tiffany Cooper, Meagan Harms, Refujia Gomez, Stephen P J Fancy, Ari J Green, Ilay Caliskan, Cathryn R. Cadwell, Bridget EL Ostrem, Mary Karalius, Lindsay Braun, Sasha Gupta, Carla Francisco, Greta Peng, Alyssa T. Reddy, Kendall Nash, Samuel J. Pleasure, Caleigh Mandel-Brehm, Thomas D. Arnold, BRUNO Consortium, Mark S. Anderson, Josiah Gerdts, Michael R. Wilson, Joseph L. DeRisi

## Abstract

Autoimmune diseases arise when B and T lymphocytes lose tolerance to self. Yet in most disorders, the underlying molecular determinants, including autoantibodies, epitopes and lymphocyte clones that drive tissue injury remain undefined. Rapid-onset obesity with hypothalamic dysfunction, hypoventilation and autonomic dysregulation (ROHHAD) is a rare and often fatal pediatric neuroendocrine syndrome with strong evidence of antigen-driven paraneoplastic autoimmunity, including association with the intracellular autoantigen ZSCAN1. However, the effector immune circuit and the epitope-level determinants operating within the hypothalamus and brainstem have remained unknown. To address this challenge in ROHHAD and more broadly in autoimmune disease, we developed the Autoimmune Epitope and immunoGlobulin/Immune-receptor identification System (AEGIS), an integrated framework that links immune repertoires to their cognate self-epitopes. AEGIS combines B cell and T cell receptor profiling from sites of tissue injury with high-resolution epitope mapping, direct sequencing of antigen-specific autoantibodies, *in silico* antibody–antigen folding, selection, and T cell antigen discovery. Applied to a deeply phenotyped child with ROHHAD, AEGIS revealed a compartmentalized, clonally restricted immune response in which brain-deposited IgG and expanded cerebrospinal fluid B cell and CD4 T cell clonotypes converged on shared ZSCAN1 epitopes, resolved to minimal determinants and peptide–MHC ligands. These findings provide a clone- and epitope-linked mechanistic map of ROHHAD autoimmunity and establish a generalizable framework for identifying candidate pathogenic clones and antigens across diverse autoimmune diseases.

---

Autoimmune diseases arise from the loss of lymphocyte tolerance to self, yet in most disorders the mechanistic chain of events linking tissue injury to the antigenic epitopes involved and the cognate B and T cell clones that drive injury remains incomplete(*1–5*).This gap persists despite major advances in protein display technologies and immune-repertoire sequencing(*6*, *7*).

Although these approaches can catalogue autoantibody reactivities and identify expanded lymphocyte clonotypes, they rarely connect these findings in a way that reveals the pathogenic cells and epitopes operating within affected tissue(*1*, *3*, *8*, *9*). As a result, the molecular mechanisms that drive disease often remain undefined, hindering the development of precise, targeted therapies(*1*, *8*).

This disconnect has important clinical consequences. Autoantibodies are widely used as diagnostic and prognostic markers across autoimmune diseases, including type 1 diabetes, systemic lupus erythematosus and paraneoplastic autoimmune encephalitides, in many cases their mechanistic contribution remains uncertain(*10–12*). The problem is especially acute for antibodies directed against intracellular antigens, which often mark disease without clearly explaining how tissue injury occurs(*11*, *13*). Even when antibodies are strongly implicated as direct effectors, mechanistic studies usually depend on serum/CSF testing and recombinant patient-derived monoclonal antibodies rather than direct analysis of antigen-bound antibodies within diseased tissue(*11*, *13*, *14*).

A parallel problem exists for T cells. Bulk and single-cell T cell receptor sequencing can reveal striking clonal expansions, often suggesting antigen-driven pathology, but the peptide–MHC ligands recognized by those T cell receptors usually remain unknown(*15*, *16*). Without methods that connect antibody and T cell clones to the self-epitopes they recognize *in situ*, the immune circuitry underlying many autoimmune diseases remains only partially defined(*4*, *15*, *17*). Together, these limitations help explain why treatment still relies largely on broad immunosuppression, which often incompletely controls disease, carries substantial toxicity, and cannot distinguish pathogenic from protective immunity(*1*, *18*, *19*).

Rapid-onset obesity with hypothalamic dysfunction, hypoventilation and autonomic dysregulation (ROHHAD) is an autoimmune disorder that exemplifies these challenges. ROHHAD is a rare and frequently lethal pediatric syndrome in which previously healthy children develop dramatic weight gain over a period of months, followed by hypothalamic dysfunction, central hypoventilation and severe autonomic instability(*20*, *21*). Existing therapies are largely empiric and often fail to prevent major morbidity or death(*20*, *22*). Several observations strongly support an antigen-driven paraneoplastic autoimmune process, including the frequent association with neural crest tumors, the subacute clinical course, and partial responses to broad immunosuppression(*22*). Autoantibodies against ZSCAN1 are associated with ROHHAD, but the effector immune cells that drive disease, and the relevant targets within the hypothalamus and brainstem, remain undefined(*21*, *22*).

To better understand complex life-threatening autoimmune disorders, such as ROHHAD, we integrated a series of technological innovations into a unified workflow called the Autoimmune Epitope and immunoGlobulin/Immune-receptor identification System (AEGIS). Here, we deploy AEGIS using important tissue, CSF, and blood sampling to construct a clone- and epitope-resolved autoimmune map defining the molecular effectors of ROHHAD. More broadly, this approach can be generalized to diverse autoimmune and paraneoplastic conditions, and may help enable more precise, patient-specific immunotherapies.

## Characterization of ZSCAN1 autoantibodies within an index ROHHAD case

A 4.5-year-old female presented to the pediatric intensive care unit with blood-pressure lability and hypoventilation. At 18 months of age she was diagnosed with neurofibromatosis type 1 (NF1), complicated by a thoracic spine plexiform neurofibroma and an optic nerve glioma that did not require therapy. At ∼3.5 years of age she developed hypothalamic–pituitary dysfunction with recurrent sodium dysregulation and was subsequently diagnosed with hypothyroidism and adrenal insufficiency of unclear etiology (**Fig. 1A**). Over the ensuing 10 months, between stabilization of her endocrinopathies and her first PICU admission, her body weight rapidly escalated from 15.09 kg (60.75th percentile) to 38.73 kg (>99.99th percentile) (**Fig. 1B**).

**Figure 1:**
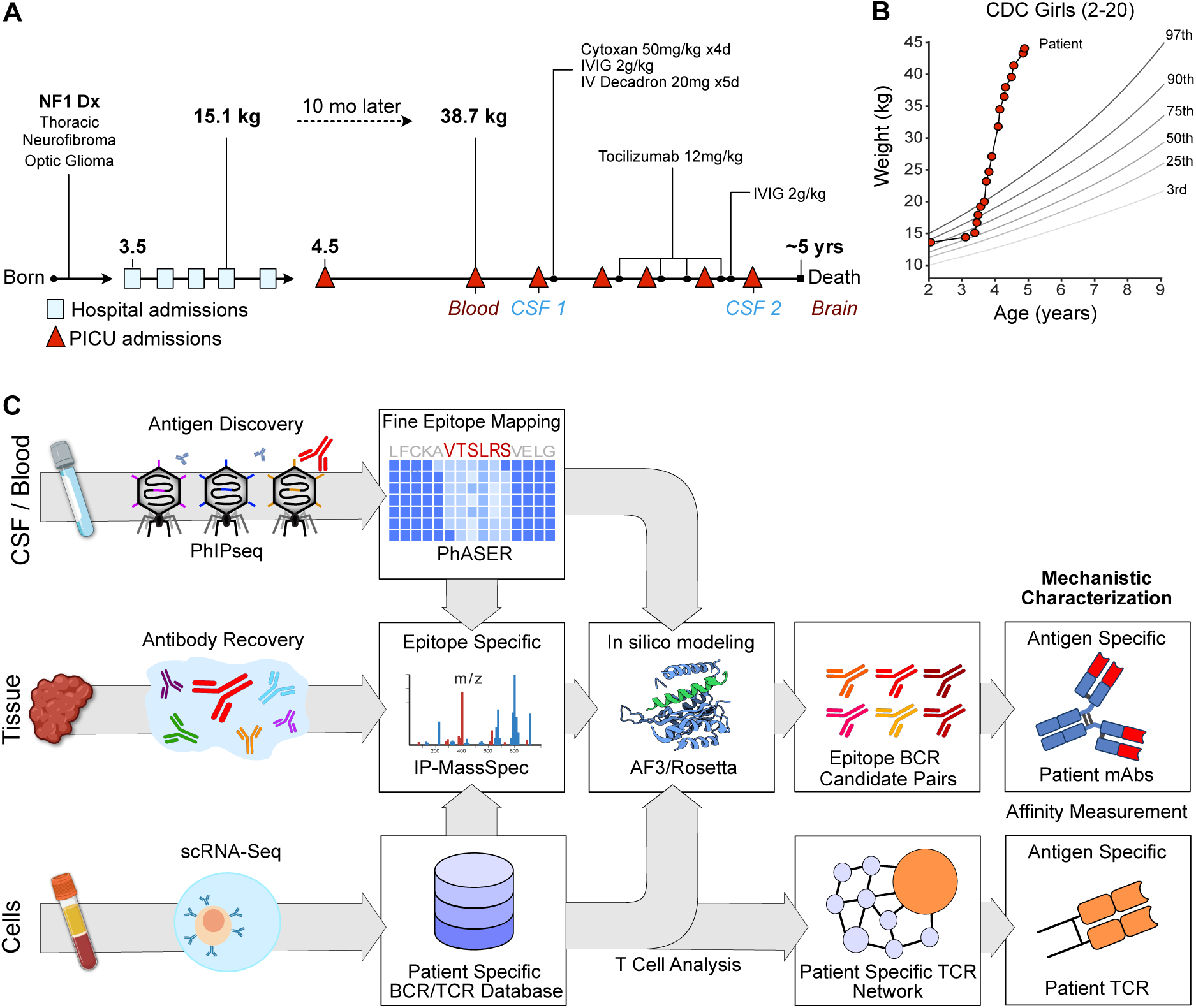
Index patient clinical course and immune profiling. **A)** Timeline depicting the disease course of the index patient. The timing of various samples acquired and utilized for experiments are shown in relationship to key clinical events and immunomodulatory therapies. **B)** The index patient’s CDC growth chart. **C)** Flow diagram of the multiple technologies comprising the AEGIS platform, and how they are integrated and applied to the index patient.

Extensive subspecialty evaluation failed to identify a unifying diagnosis. Given the subacute onset and constellation of hypothalamic, respiratory, and autonomic features in a previously stable child, an autoimmune hypothalamic encephalopathy was suspected, and she was enrolled in the UCSF Neuroinflammatory Disease (IRB #13-12236) cohort and the UCSF Child Health Advanced Molecular Phenotyping (CHAMP) program for deep immunological profiling **(Fig. 1C)**.

The AEGIS workflow is summarized in Figure 1C. Antibody profiling from serum or CSF is a practical entry point and proteome-wide screening technologies enable unbiased discovery at scale(*7*).The index patient’s serum and CSF was interrogated using a previously validated and customized PhIPseq human library (∼730,000 clones) featuring tiled 49 amino acid segments of the human proteome(*21*, *23–27*).PhIP-Seq was performed by incubating serum or CSF with the library and conducting sequential rounds of immunoprecipitation to enrich antibody-bound peptides(*28*);enrichment was quantified as fold-change over mock immunoprecipitation (FC over mock-IP) and compared against serum from 48 healthy adult donors. Consistent with previous results from this lab and others, ZSCAN1 emerged as the dominant target, with multiple ZSCAN1-derived peptides ranking among the top five enriched peptides in both CSF timepoint 1 and CSF timepoint 2, and with striking aggregate enrichment (sum of canonical-isoform peptides) in CSF (FC over mock-IP: 31,831 at timepoint 1; 8,772 at timepoint 2)(*21*, *29*, *30*).Highlighting the importance of interrogating tissues or fluids anatomically associated with the disease, ZSCAN1 peptides were also detectable in serum but at substantially lower levels (aggregate FC over mock-IP: 239). Although aggregate ZSCAN1 enrichment in index serum exceeded all 48 healthy control sera, some enrichment was observed in a small subset of controls (3/48), consistent with prior reports (**Fig. 2A**)(*21*).Despite aggressive therapy with intravenous (IV) immunoglobulin (IG), IV cyclophosphamide, high-dose IV steroids and IV tocilizumab, the patient’s condition progressed, marked by increasingly frequent PICU admissions. She died at home approximately 4 months after initiation of immunomodulation from profound nocturnal hypoventilation leading to cardiac arrest.

**Figure 2:**
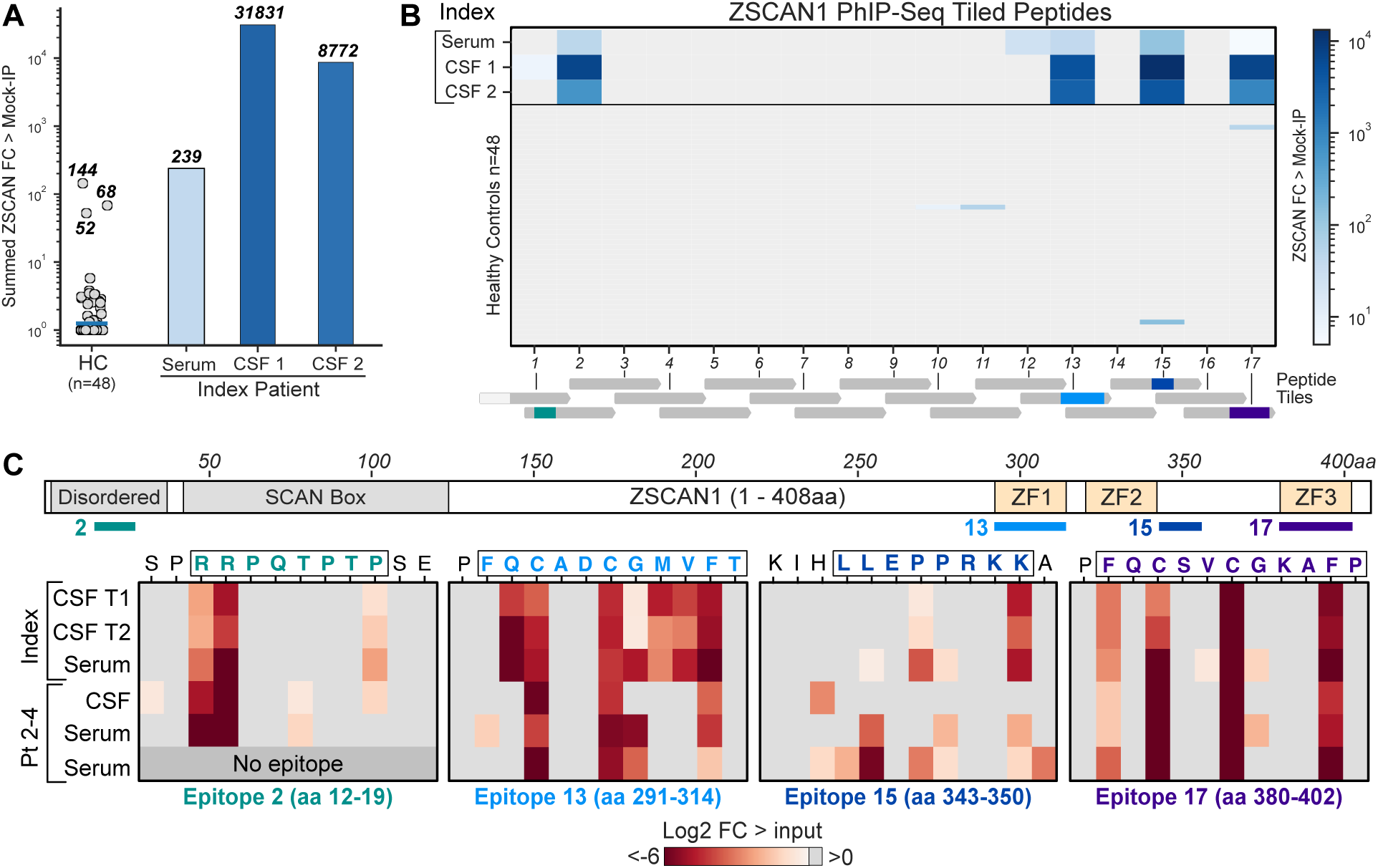
Distribution of ZSCAN1 epitopes across ROHHAD patients and healthy controls. **A)** ZSCAN1 signal (fold change over mock-IP) was quantified by summing enrichment values across all library peptides mapping to the canonical 408 amino acid ZSCAN1 protein. Healthy control (HC) samples are shown as individual jittered points, with the HC median indicated by a horizontal line. Index patient samples are shown as bars for serum, CSF timepoint 1 (#1) and CSF timepoint 2 (#2). The y-axis is on a logarithmic scale; for visualization, all values ≤1 were floored to 1 prior to plotting**. B)** Heatmap showing distribution of ZSCAN1 signal (fold change over mock-IP) across each ZSCAN1 PhIP-Seq fragment in healthy controls (n=48) and index patient serum, CSF #1, and CSF #2. **C)** Results of PhASER epitope mapping “single” scan for each of the 4 most reactive peptides identified by PhIP-Seq. Complete epitopes 2 and 15 are shown; only the N-terminal portion of epitopes 13 and 17 fit into this visualization, but the full epitope coordinates are described and mapped onto the ZSCAN1 schematic. Signal is calculated as log2 fold-change over the input library. Top half maps the epitopes for the index patient in serum, CSF #1, and CSF #2. Bottom half maps the epitopes across 3 additional ROHHAD patients.

ZSCAN1 is a 408 amino acid nuclear transcription factor containing a SCAN domain and three cysteine-2 histidine-2 (C2H2) zinc-finger (ZF) motifs. ZSCAN1 expression is highly restricted, with predominant expression in the epididymis and central nervous system(*31*, *32*). Within the CNS, ZSCAN1 expression is highest in the hypothalamus and midbrain(*31*, *32*).Across index samples, ZSCAN1 reactivity spanned multiple distinct regions of the protein, including sequences within the zinc-finger domains. In contrast, each of the ZSCAN1-reactive healthy outliers exhibited enrichment restricted to a single region. Two ZSCAN1 peptide fragments from our PhIP-Seq library (2 and 13) were enriched in the index case but were absent in healthy controls (**Fig. 2B**).

To precisely delineate the autoreactive ZSCAN1 epitopes, we employed a natural extension of our previously published deep scanning mutagenesis PhIP-Seq system(*27*).**Ph**age-**A**ssisted **S**canning **E**pitope **R**ecovery (PhASER) is a highly parallel, high-resolution epitope-mapping system with 3 key design elements. First, to identify critical binding residues, within each PhIP-Seq-positive 49-mer, every amino acid is individually mutated to alanine (or to glycine if alanine was the native residue). Loss of antibody binding to any single-residue mutant defined that position as essential for recognition. Second, to define the exact C-terminal limit of the epitope, each codon was sequentially replaced with a stop codon. Antibody binding occurs only when the displayed fragment fully contains the epitope, enabling precise determination of the C-terminal border. Finally, to identify the N-terminal limit, we progressively extended an N-terminal XTEN linker—chosen for its non-immunogenicity and minimal structural interference—one residue at a time. Binding abruptly disappears once the linker encroaches into the epitope, revealing the N-terminal boundary **(Extended Data Fig. 1)**.

Using a custom phage library containing ZSCAN1 PhASER tiles, four distinct ZSCAN1 epitopes were mapped in the index patient. Epitope 2 (aa 12–19; within ZSCAN1 peptide fragment 2) localized to the disordered N-terminus. Epitope 13 (aa 292–314; within ZSCAN1 peptide fragment 13) exactly matched ZF1 (aa 292–314). Epitope 15 (aa 343–350; within ZSCAN1 peptide fragment 15) began one residue C-terminal to ZF2 (aa 320–342). Epitope 17 (aa 380–402) precisely matched ZF3 (aa 380–402; within ZSCAN1 peptide fragment 17) (Figure 2C, complete maps in **Extended Data Fig. 2**). Epitopes 2 and 13 fell within regions that were absent from all 48 healthy controls. All four epitopes were detectable with identical residue-level specificity in both serum and CSF from the index case, and remained unchanged in CSF obtained approximately three months.

PhASER was applied to three additional ZSCAN1 antibody positive ROHHAD patients (one CSF sample and two serum samples). Each additional ROHHAD patient sample targeted at least three of the four epitopes and shared highly similar critical-residue patterns across epitopes, indicating convergent fine specificity across independent cases (**Fig. 2C, bottom**). Together, these data support the notion that a multi-epitope autoreactive response to ZSCAN1 is stereotypical amongst patients.

## Expanded CNS B cell clonotypes encode ZSCAN1-specific receptors

To characterize the autoimmune drivers of ROHHAD at the cellular level, immune cells from the CSF of the index patient were characterized by single cell sequencing. Across 29 CSF B cells (inclusive of both timepoints), 14 unique BCR clonotypes were recovered. To broaden candidate discovery, BCRs were also sequenced from matched PBMCs, identifying 204 peripheral B cells (200 clonotypes). Because antigen-experienced, class-switched IgG B cells are thought to be particularly relevant for CNS antigen presentation, subsequent analyses focused on the 81 IgG isotype BCRs (**Fig. 3A**).

**Figure 3:**
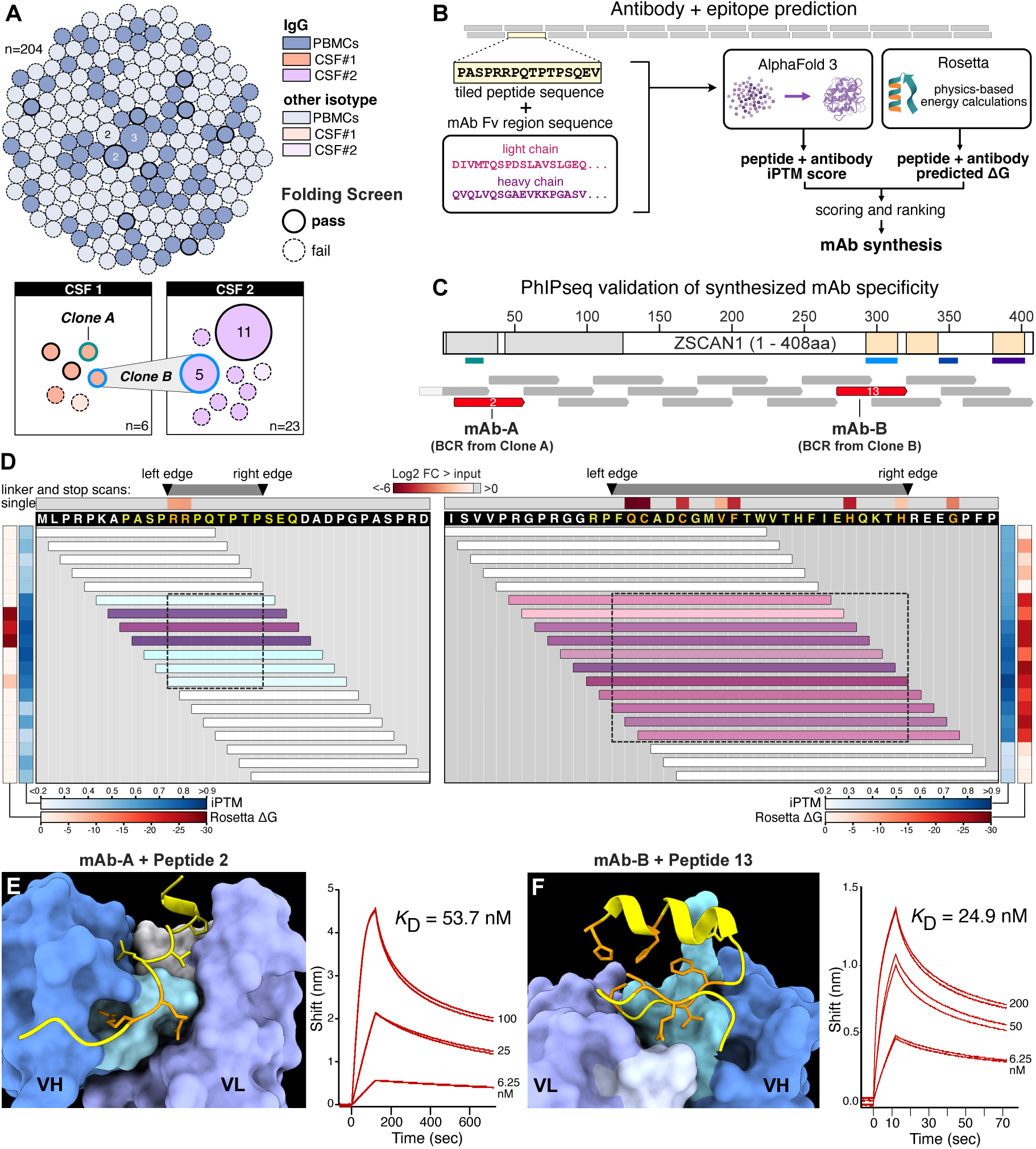
Identification of distinct CNS B cell lineages binding ZSCAN1 with high affinity and specificity using the PAIRIS pipeline. **A)** Clonal repertoire of B cells sampled from blood (PBMCs) and two CSF timepoints (CSF#1 and CSF#2). Each circle represents a unique BCR clone, with size proportional to clonal frequency; n indicates the total number of cells sampled per compartment. Solid outlines denote clones predicted to bind to at least one ZSCAN1 peptide by PAIRIS (iPTM >= 0.85 and negative Rosetta binding energy; while dashed outlines indicate clones not predicted to bind ZSCAN1. **B)** PAIRIS (Prediction of Antibody-antigen Interaction at high Resolution In Silico) workflow for high-throughput binding prediction. Overlapping peptides (15mer + 25mer) spanning ZSCAN1 and monoclonal antibody variable region (mAb Fv) sequences are input to AlphaFold3 for structure prediction of peptide-antibody complexes. Predicted structures are evaluated by interface predicted template modeling (iPTM) scores and Rosetta-calculated binding energies (ΔG) to identify binding interactions. **C)** Monoclonal antibodies from clones A and B identify distinct, non-overlapping ZSCAN1 peptide fragments 2 and 12, respectively (aa 273-321 and 9-57; UniProt: Q8NBB4). **D)** Validation of PAIRIS binding predictions by PhASER and biolayer interferometry (BLI). Vertical column heatmaps display iPTM scores (blues, average interface confidence between peptide and both antibody chains) and Rosetta ΔG values (reds) for mAb-A against 15-mer peptides spanning ZSCAN1 aa 1-33 (left) and mAb-B against 25-mer peptides spanning aa 279-321 (right) (UniProt: Q8NBB4). Horizontal bars represent peptides input into PAIRIS and span the indicated amino acid positions. The color of these bars represent the sum of iPTM and Rosetta values, with deeper purple indicating more favorable iPTM and Rosetta ΔG values. Above the amino acid sequence, the horizontal row heatmap shows the PhASER “single” scan results for mAb-A (left) and mAb-B (right), with critical residues highlighted in orange in the sequence. Residues highlighted in yellow are part of the peptide shown in E,F. The “left edge” and “right edge” arrows represent the edges of the epitope as determined by PhASER “linker” and “stop” scan results (**Extended Data** Fig. 4). Dashed rectangles denote PhASER epitope regions across colored horizontal bars. **E, F)** AlphaFold3-predicted structures show the highest-scoring peptide-antibody complexes with critical epitope residues highlighted (orange). BLI sensorgrams using peptides that include the respective epitope region confirm binding with calculated KD values.

Experimentally screening large numbers of candidate BCR sequences for antigen specificity is slow, expensive, and often impractical when patient-derived repertoires contain many plausible clones. To overcome this bottleneck, we developed Prediction of Antibody–antigen Interaction at high Resolution In Silico (PAIRIS), a computational framework designed to prioritize likely BCR–antigen interaction candidates before experimental testing. PAIRIS uses AlphaFold3 to model patient-derived IgG Fabs in complex with peptides spanning the full coding sequence of a candidate target protein, using 15- and 25-mers sliding-window tiles generated at single-amino-acid resolution (**Fig 3B**)(*33*). By systematically evaluating each predicted antibody–peptide complex, PAIRIS enables rapid, high-resolution triage of large candidate repertoires that would otherwise be prohibitively difficult to test empirically. For each antibody–peptide pair, we assessed the predicted complex using the interface predicted template modeling (iPTM) score, for which values greater than 0.8 are generally considered indicative of a high-confidence interaction(*34*). In parallel, we used Rosetta to compute a binding energy score for each modeled complex, with more negative values indicating more favorable predicted binding (Methods)(*35*). Using conservative thresholds (iPTM >0.85 and negative Rosetta binding energy), PAIRIS predicted 14 of 81 IgG BCRs to bind at least one ZSCAN1 peptide **(Fig. 3B)**. Predicted binders were enriched in CSF: 5 of 14 CSF IgG clonotypes met these criteria, including the only clonotype that persisted and expanded across time points (Clone B) (**Fig. 3A**).

Monoclonal antibodies were subsequently synthesized and expressed corresponding to the five CSF clonotypes prioritized by PAIRIS (mAb-A from Clone A, mAb-B from Clone B, etc.) Each mAb was evaluated by independent PhIP-Seq experiments against the full human peptidome to determine specificity. Three of the five antibodies—mAb-A, mAb-B, and mAb-C—selectively enriched different ZSCAN1-derived peptides, each with fragment-level specificity, indicating distinct ZSCAN1 reactivities across intrathecal lineages (**Fig. 3C**). Whereas mAb-A and mAb-B preferentially enriched fragments detected in bulk patient serum or CSF (ZSCAN1 peptide fragments 2 and 13, respectively), mAb-C enriched peptide fragment 5 (SCAN-box region), despite no detectable enrichment of this fragment in serum or CSF at either time point. (**Fig. 3C**). Phylogenetic analysis indicated that each ZSCAN1-reactive BCR arose from distinct germline rearrangements (**Extended Data Fig. 3**).

To define epitope specificity at single-amino acid resolution, mAb-A and mAb-B were mapped using a custom ZSCAN1 PhASER library revealing that mAb-B bound Epitope 13 (aa 292–314; ZF1) and mAb-A bound Epitope 2 (aa 12–19; disordered N-terminus), matching the epitopes independently mapped in index serum and CSF and notably corresponding to the two epitope regions absent in all healthy controls (**Fig. 3D, Fig. S4**). Because ZSCAN1 peptide fragment 5 was not enriched in bulk patient samples, it was not included in the initial PhASER design. For the mapped antibodies, refined *in silico* complex models recapitulated the PhASER-defined critical contact residues, supporting concordance between PAIRIS predictions and experimental epitope definition (**Fig. 3D**). To identify which residues of ZSCAN1 peptide fragment 13 are most critical for binding mAb-B, we computed the per-residue change in solvent-accessible surface area (delta SASA) between the unbound peptide and antibody-bound complex. Delta SASA values, together with heavy/light chain contact distances to the peptide, identified the key contact residues mediating mAb-B–ZSCAN1 peptide fragment 13 binding. Consistent with these calculations, in silico deep mutational scanning of the peptide confirmed that residues most critical for complex stability correspond with PhASER-defined critical contact residues (**Extended Data Fig. 5**). PAIRIS modeling was also performed for mAb-C and ZSCAN1 peptide fragment 5, providing predicted epitope mapping in the absence of PhASER data (**Extended Data Fig. 9**).

To orthogonally validate specificity and quantify binding, the minimal PhASER-defined epitopes were analyzed by bio-layer interferometry (BLI). mAb-B and mAb-A bound their respective ZSCAN1 peptides with affinities of 24.9 nM and 53.7 nM, respectively (**Fig. 3D**). mAb-C also bound the ZSCAN1 peptide fragment 5, although reliable affinity estimation was limited by peptide insolubility (**Extended Data Fig. 9**). Collectively, these data link expanded CSF B cell clonotypes to ZSCAN1 epitopes at single-amino-acid resolution and confirm the utility of PAIRIS as a scalable framework for prioritizing candidate antigen specificities for experimental validation.

Matched single-cell transcriptomes were available for 5 ZSCAN1-reactive B cells, including 2 from timepoint 1 and 3 from timepoint 2. These cells mapped predominantly to memory and intermediate B-cell states, with 4 of 5 cells assigned to these populations. All 4 expressed high levels of HLA-DRA and HLA-DRB1, consistent with active MHC class II antigen presentation. Clone B (mAb-B lineage) persisted to timepoint 2 and expanded from 1 to 5 cells, becoming the second most expanded clone in the CSF. Notably, all 3 ZSCAN1-reactive cells with matched transcriptomes at timepoint 2 derived from this clone and spanned intermediate B-cell, memory B-cell and plasmablast states. The memory B cell expressed CD69, CD80 and CD86, consistent with an activated, T cell-interacting state, whereas the plasmablast showed high IGHG1 expression, consistent with active IgG1 production. Collectively, these findings support ongoing activation and differentiation of ZSCAN1-reactive B cells in the CSF, including a state compatible with antigen presentation to CD4+ T cells (**Extended Data Fig. 5**).

## Brain-deposited IgG is clonally restricted and regionally structured

Neuropathologic evaluation of autopsy specimens revealed a diffuse inflammatory process, including expansion of the leptomeningeal spaces with predominantly macrophage and T cell infiltrates. No evidence of infection was identified. Reactive astrogliosis and focal microglial activation were present in multiple brain regions, including the hypothalamus and brainstem (**Extended Data Fig. 7**). Despite limited lymphocyte infiltration into the brainstem at the time of death, post-mortem brainstem sections showed marked IgG deposition compared with an age-matched control (**Fig. 4A**). To determine whether this reflected antigen-specific binding within tissue rather than passive diffusion from blood or CSF, IgG was first eluted from fresh brain sections, then profiled using PhIP-Seq.

**Figure 4:**
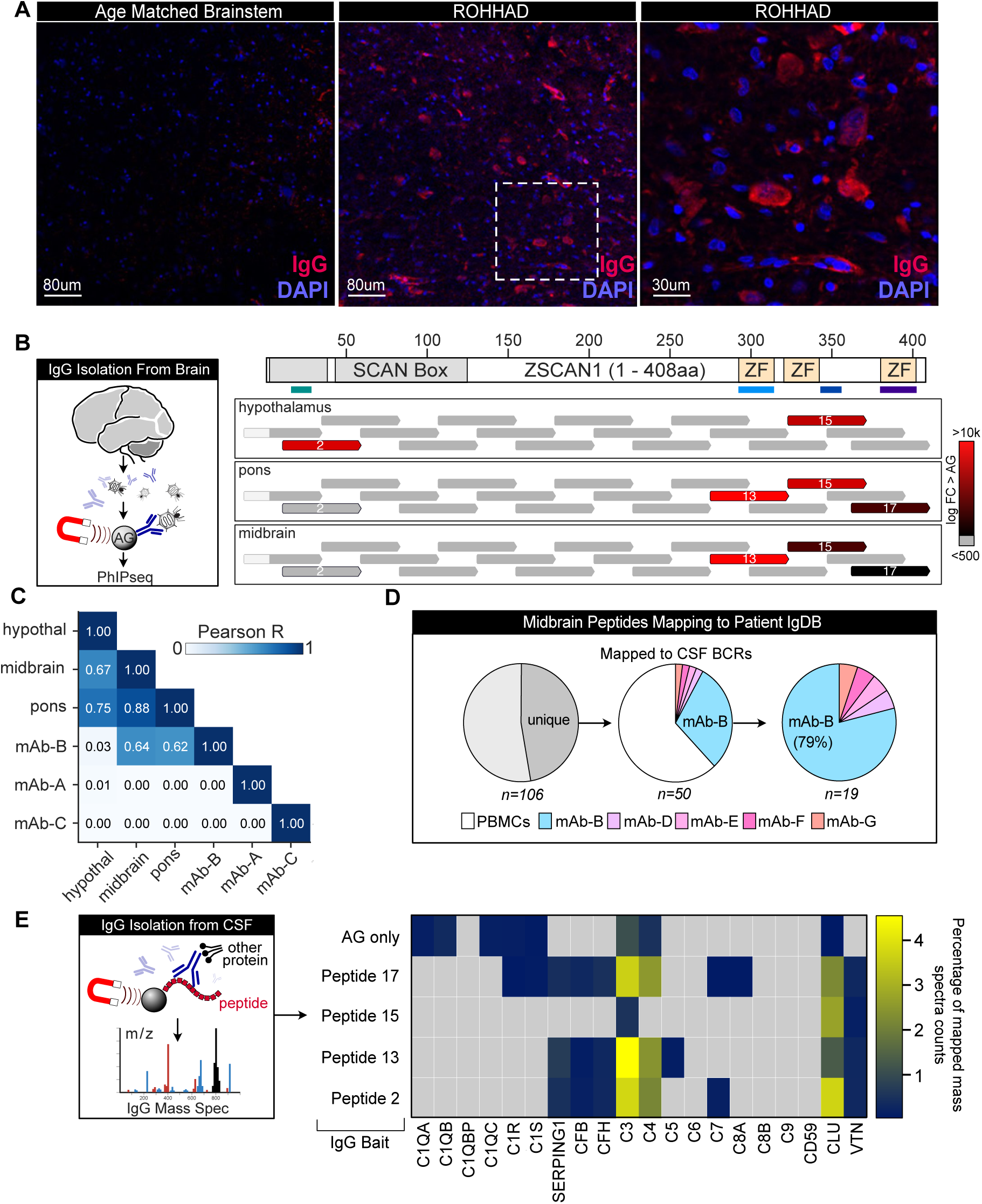
ZSCAN1 autoantibody deposition within brain tissue. **A)** Immunofluorescence staining of post-mortem brainstem showing increased IgG deposition (red) in the ROHHAD case compared with an age-matched control; nuclei are counterstained with DAPI (blue). Representative low- and high-magnification (zoom in of the white hashed rectangle) fields are shown. Scale bars, 80 µm (left and middle) and 30 µm (right). **B)** Workflow for regional elution of tissue-deposited IgG followed by PhIP–Seq profiling, and mapping of enriched peptides along ZSCAN1 (aa 1–408; UniProt: Q8NBB4). Tracks show relative enrichment of ZSCAN1 fragments across hypothalamus, pons and midbrain eluates, highlighting regional structure (hypothalamus dominated by fragment 2; brainstem regions preferentially enriching fragment 13). Color indicates peptide enrichment magnitude (fold-change). **C)** Pairwise similarity of PhIP-Seq enrichment profiles across brain regions and CSF-derived ZSCAN1-reactive monoclonal antibodies (mAb-A, mAb-B, mAb-C), quantified by Pearson correlation (r). **D)** Antibody repertoire mass spectrometry (Ig-MS) strategy and results linking tissue-deposited IgG to intrathecal B cell clonotypes. IgG immunoprecipitated from CSF (timepoints 1 and 2) and midbrain was digested and peptide spectra were mapped to a patient-specific BCR reference database generated from single-cell BCR sequencing. Pie charts summarize uniquely mapping peptides and their assignment to PBMC- versus CSF-derived BCR sequences, and the distribution of CSF-derived assignments across clonotypes, demonstrating marked enrichment of clone B/mAb-B in midbrain. **E)** Epitope-directed immunoprecipitation followed by mass spectrometry (IP-MS) using PhASER-defined minimal epitopes to capture ZSCAN1-specific IgG from CSF. Heatmap summarizes co-precipitating proteins across capture conditions, highlighting preferential enrichment of complement components (including C3 and C4) in mAb-B (binding ZSCAN1 peptide fragment 13) epitope-directed immunoprecipitates compared with protein A/G capture.

Relative to serum and CSF, the brain-deposited autoreactive repertoire was markedly restricted. Using a fold-change enrichment cutoff of greater than 10, there were 932 autoreactive peptides within serum, 383 in terminal CSF (timepoint 2), and fewer in brain regions (305 in hypothalamus, 142 in pons, 134 in midbrain) (**Extended Data Fig. 8**). Interestingly, CNS regions exhibited distinct enrichment patterns. Hypothalamic eluates were dominated by ZSCAN1 peptide fragment 2 (followed by fragment 15), whereas brainstem regions (midbrain and pons) preferentially enriched peptide fragment 13 (**Fig. 4B**). In contrast, serum and CSF enriched ZSCAN1 peptide fragments 2, 13, and 15 more evenly, suggesting that tissue deposition is selective rather than a simple reflection of circulating antibodies. In addition to ZSCAN1, brain eluates were enriched for other zinc-finger proteins, including ZNF616 and ZNF836, which were among the top 10 most enriched peptides in each brain section.

To link tissue-deposited antibodies to intrathecal B cell clonotypes, regional PhIP-Seq profiles were compared with CSF-derived mAbs with confirmed ZSCAN1 reactivity (mAb-A, mAb-B, mAb-C). Midbrain and pons profiles were highly concordant (r = 0.880), and both were most similar to mAb-B (midbrain r = 0.644; pons r = 0.622) (**Fig. 4C**). mAb-B recapitulated key features of the brainstem signature, including preferential enrichment of ZSCAN1 peptide fragment 13 and ZNF616/ZNF836 peptides. Consistent with this, PAIRIS modeling predicted favorable interactions between mAb-B and ZNF616/ZNF836 peptides containing the Zn finger motifs, which we confirmed by BLI: mAb-B bound ZNF616 and ZNF836 with high affinity (KD 6.58 nM and 12.1 nM), exceeding its affinity for the ZSCAN1 peptide epitope (**Extended Data Fig. 6**). Given the extensive somatic hypermutation present in mAb-B (10.2%) relative to mAb-A (0.6%), these data raise the possibility that affinity maturation may resulted in an expanded binding spectrum including structurally related zinc-finger motifs.

Finally, enrichment of mAb-B in brain was validated using antibody repertoire mass spectrometry (Methods). IgG immunoprecipitated from both CSF time points and midbrain was digested and mapped to a patient-specific BCR reference database generated from single-cell BCR sequencing. Because antibody variable regions contain substantial shared sequence, we focused on peptides that mapped uniquely to a single BCR sequence. At both CSF time points, only ∼10% of uniquely mapping peptides corresponded to CSF-derived BCR sequences (rather than the PBMC database). In contrast, the midbrain showed substantially greater enrichment: 38% of uniquely mapping peptides originated from the CSF database, and the majority mapped to BCR clone B/mAb-B **(Fig. 4D).**

To further assess clonotypic specificity in tissue, we applied a CDR3-based identity filter (>50% heavy- or light-chain CDR3 coverage). In CSF, 44 antibodies (timepoint 1) and 124 antibodies (time point 2) met this criterion, consistent with a polyclonal intrathecal repertoire. In contrast, in midbrain eluates only two antibodies passed this filter, and only mAb-B passed both the heavy and light chain filters, suggesting marked clonal restriction among brain-deposited IgG. Across samples, Ig-MS provided deep coverage of the mAb-B variable regions (∼90% of the heavy-chain variable region and ∼98% of the light-chain variable region) **(Extended Data Fig. 10)**. Together, these data indicate that CNS-deposited antibodies form a narrowed, regionally structured repertoire dominated in the brainstem by one expanded intrathecal lineage (Clone B/mAb-B).

## ZSCAN1-specific antibodies co-purify with complement

Autoantibodies to intracellular antigens are often treated as non-pathogenic biomarkers; however, in related paraneoplastic neurologic syndromes, experimental models support antibody-linked effector mechanisms that may amplify tissue injury(*36*, *37*). Given the selective deposition of ZSCAN1- and zinc-finger-reactive IgG within diseased brain, immunoprecipitation followed by mass spectrometry (IP-MS) was used to detect associated proteins, especially complement factors. Using PhASER-defined minimal epitopes as bait, epitope-specific IgG was immunoprecipitated from CSF and analyzed for co-precipitating proteins by MS. Immunoglobulin peptide mapping re-validated the expected specificities, with mAb-B enriching ZSCAN1 peptide fragment 13 and mAb-A enriching ZSCAN1 peptide fragment 2, whereas protein A/G immunoprecipitation recovered all three ZSCAN1 antibodies more evenly.

Strikingly, complement components C3 and C4 were among the most enriched co-purifying proteins in the mAb-B epitope-directed immunoprecipitates, suggesting these IgG1 brain-deposited antibodies are complement associated in CSF, and supporting a complement-coupled effector program at sites of CNS injury (**Fig. 4E**).

## Expanded CSF CD4 T cell clonotypes recognize ZSCAN1-derived peptide–MHC ligands

The presence of clonally expanded, somatically hypermutated B cells in the CSF targeting a shared autoantigen suggests coordinated T cell–B cell immunity within the central nervous system. Such B cell responses typically require cognate CD4 T cell help during priming or reactivation, and their persistence within the intrathecal compartment suggests ongoing or repeated antigen encounter rather than passive diffusion from the periphery. Moreover, because ZSCAN1 is an intracellular protein, antibody recognition alone is unlikely to fully explain immune activation or tissue inflammation, whereas CD4 T cells recognizing processed ZSCAN1-derived peptides could provide both helper signals to B cells and a direct source of pro-inflammatory effector activity within the CNS. Together, these observations led us to hypothesize that CD4 T cells recognizing ZSCAN1-derived peptide–MHC complexes could play a role in ROHHAD pathogenesis.

To assess circulating ZSCAN1-reactive T cells, we incubated patient PBMCs with an overlapping ZSCAN1 peptide pool (15-mers with 11-aa overlap across full-length ZSCAN1) and quantified activation using an activation-induced marker (AIM) assay (CD137, OX40, CD69)(*38*). ZSCAN1 stimulation activated 0.1 percent of circulating T cells, indicating that ZSCAN1-reactive T cells in blood, if present, were at very low frequency (**Extended Data Fig. 11**).

CSF was prioritized as the most relevant compartment and single-cell RNA sequencing was performed with paired TCR sequencing at two time points separated by ∼3 months. Across both timepoints, we identified 481 unique paired αβ TCRs among 601 T cells. At timepoint 2, 16 expanded CD4 clonotypes were detected, 9 of which persisted across both timepoints (**Fig. 5A**).

**Figure 5:**
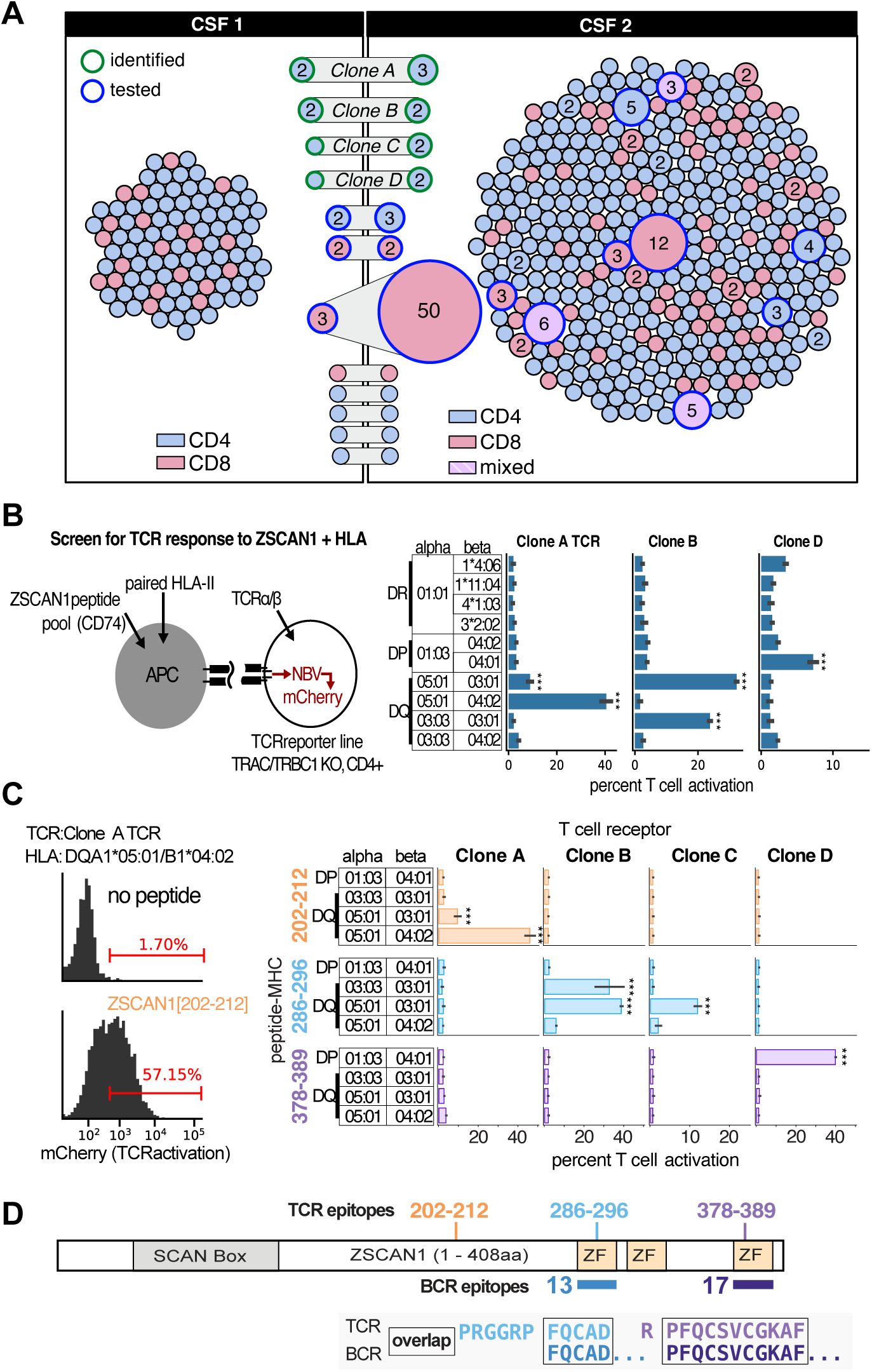
Expanded intrathecal CD4 T clonotypes are ZSCAN1-specific and track with B cell epitope usage. **A)** Single-cell paired αβ TCR repertoire structure in CSF collected at two time points (∼3 months apart). Each circle represents a unique T cell clonotype; clonotype size reflects abundance and colors denote lineage assignment (CD4 versus CD8, as indicated). Expanded CD4 clonotypes detected at time point 2 and persistent clones shared across time points are highlighted. A dominant CD8 clonotype present at both time points is indicated (numbers within highlighted nodes denote clone size). **B)** Functional screening of expanded CSF CD4 TCR clonotypes. TCRs were expressed in TCR-deficient Jurkat NFAT reporter cells co-expressing CD4 and co-cultured with APCs expressing individual HLA class II allele pairs and a pooled library of ZSCAN1 peptides fused to CD74. Reporter activation (mCherry) quantified responses across HLA-DR, -DP, and -DQ allele pairs. **C)** Identification of minimal ZSCAN1 peptide antigens and HLA restriction. Left, representative flow cytometry plots showing activation of clone A reporter cells in response to ZSCAN1[202–212] presented by HLA-DQA1*05:01/DQB1*04:02. Right, candidate peptides were tested across the indicated HLA class II allele pairs, defining minimal activating peptides and restricting HLA contexts: clone A recognized residues 202–212 in the context of HLA-DQ (DQA1*05:01 paired with DQB1*04:02 or DQB1*03:01), clones B and C recognized residues 286–296 in the context of HLA-DQ (DQA1*03:03 or DQA1*05:01 paired with DQB1*03:01), and clone D recognized residues 378–389 in the context of HLA-DP (DPA1*01:03/DPB1*04:01). **D)** Integration of validated T cell epitopes with ZSCAN1 antigen architecture and intrathecal B cell epitope usage. Peptide coverage across full-length ZSCAN1 (amino acids 1–408; UniProt Q8NBB4) is shown with annotated SCAN box and zinc-finger domains, validated T cell epitopes, and previously defined intrathecal B cell epitopes. ***P < 0.001 by paired one-tailed t test.

To evaluate whether expanded CSF CD4 T cell clonotypes recognized ZSCAN1-derived peptide–MHC complexes, we expressed patient-derived TCRs in a reporter cell system assessed antigen-specific activation in the presence of antigen-presenting cells expressing relevant HLA class II molecules(*39*). Antigen presentation was achieved using a targeted peptide delivery approach spanning the ZSCAN1 protein, and T cell activation was quantified by reporter readout (*40*). For three clonotypes (A, B, D), reporter activation was observed with specific HLA class II allele pairs, consistent with HLA-restricted recognition of ZSCAN1-derived peptides (**Fig. 5B**).

To define cognate epitopes, candidate peptide regions were evaluated in the context of the identified HLA restrictions, enabling identification of minimal activating sequences for each TCR. Clone A recognized residues 202–212 in the context of DQA1*05:01 paired with DQB1*04:02 or DQB1*03:01, clone B recognized residues 286–296 in the context of DQA1*03:03 or DQA1*05:01 paired with DQB1*03:01, clone C recognized the same region as clone B in a more restricted context of DQA1*05:01/DQB1*03:01, and clone D recognized residues 378–389 in the context of HLA-DPA1*01:03/DPB1*04:01 **(Fig. 5C).** The identified ligands mapped to three discrete regions of the ZSCAN1 protein, and two of these regions overlapped antibody epitopes independently defined by PhASER, indicating convergent B cell and CD4 T cell targeting of shared regions within the same intracellular autoantigen (**Fig. 5D**).

Single-cell transcriptional profiles indicated these antigen-specific clones were primarily CD4 effector memory cells, with Th1 and Th17 programming. Notably, several of these ZSCAN1 reactive CD4 cells had transcriptional signatures typical of T follicular helper (Tfh) cell, consistent with functioning within a tertiary lymphoid structure. Differential gene expression analysis of ZSCAN1-reactive versus other T cells identified GPR25 as the top upregulated gene and CD55 as the top downregulated gene, consistent with a tissue-resident effector state in an inflamed, complement-active niche **(Extended Data Fig. 12).** Together, these data show that clonally expanded T cells and B cells in ROHHAD converge on a shared specificity for ZSCAN1.

## Discussion

ROHHAD is a rare but devastating childhood syndrome with strong evidence for paraneoplastic autoimmunity, most often in association with neural crest tumors. Although its clinical phenotype is distinctive, the underlying immune architecture may apply more broadly to tissue-specific autoimmune disease, particularly to paraneoplastic neurologic syndromes marked by autoantibodies against intracellular antigens. Here, we have deeply investigated a fatal case of ROHHAD to better understand the specific elements of both the B cell and T cell axes, in the periphery, in the CSF, and in the affected tissue of the brain. In doing so, we provide direct experimental support for models advanced over decades by Darnell and others, in which B and T cell responses converge on a shared intracellular antigen, but which have rarely been resolved with sufficient precision to demonstrate this definitively(*41*, *42*). Most strikingly, ZSCAN1-reactive B and T cell clones were not merely detectable within the CNS; they were among the most expanded and persistent clones identified, arguing for particular biological importance. Moreover, these responses converged not only on the same antigen, but on shared epitopes, a finding whose mechanistic significance remains to be determined.

The initiating event in ROHHAD is likely paraneoplastic, although the precise trigger in this patient remains unknown. The pre-existing diagnosis of NF1 raises the possibility that aberrant tumor-associated expression of neural antigens, including ZSCAN1, may have contributed to the initial break in tolerance. Regardless of the initiating insult, our data support a model in which disease is sustained by an antigen-driven, self-reinforcing immune circuit within the central nervous system. In this framework, the ZSCAN1-reactive B–CD4 T cell axis functions as the antigen-specific amplifier of CNS autoimmunity. The proximate effectors of neuronal injury may include a broad range of inflammatory mechanisms, but durable pathology is likely to depend on a local circuit capable of capturing released antigen and converting it into sustained immune activation. ZSCAN1-reactive B cells are well suited for this role: following neuronal injury, they can acquire extracellular ZSCAN1 through the B cell receptor, process it, and present ZSCAN1-derived peptides to cognate CD4 T cells, thereby maintaining a focused helper response within the CNS. CD4 T cell help would in turn reinforce the autoreactive B cell compartment, promote plasma cell differentiation and continued ZSCAN1 autoantibody production, and sustain an inflammatory milieu permissive for ongoing cytotoxic activity. In this setting, the downstream effectors need not themselves recognize ZSCAN1 directly, provided they target neurons within the same injured tissue environment. Neuronal destruction would then release additional intracellular antigen, fueling renewed B cell capture and presentation. The result is a self-reinforcing loop in which tissue injury drives antigen release, antigen release drives cognate B–T cell collaboration, and cognate B–T cell collaboration sustains further tissue injury **(Fig. 6)**.

**Fig. 6.**
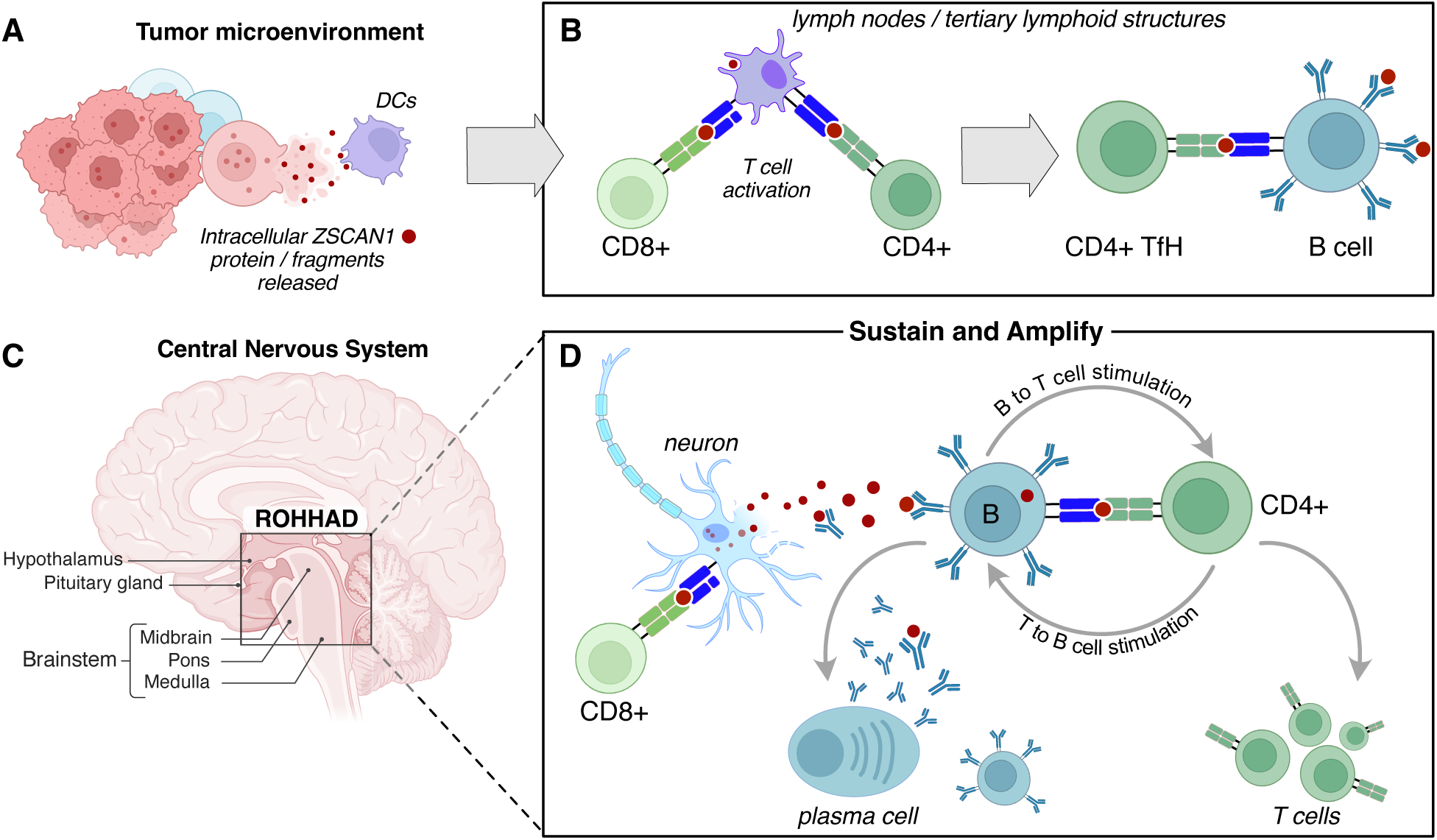
Model of a ZSCAN1-reactive immune circuit driving ROHHAD CNS autoimmunity. **A)** The initiating trigger may be aberrant expression of ZSCAN1 within a neural crest tumor in the periphery with subsequent destruction of ZSCAN1 expressing tumor cells and release of the ZSCAN protein/fragments. **B)** ZSCAN1 antigen may be filtered to lymph nodes or lymphoid structures and presented to B and T cells,leading to the development of ZSCAN1 reactive lymphocytes which subsequently traffic into the CNS. **C)** ROHHAD involves pathology to the hypothalmus, and the anatomically adjacent brainstem, which may be particularly vulnerable to autoreactive lymphocytes due to its neuroendocrine functions requiring specialized neurovascular and CSF-facing interactions. **D)** Once within the hypothalamus/brainstem, CD8 T cells may serve as direct effectors of neuronal injury. Killing of neurons containing ZSCAN1 releases intracellular antigen into the surrounding CNS microenvironment. In this framework, the pathogenic CD8 response does not require direct recognition of a ZSCAN1-derived peptide, so long as the targeted neurons express ZSCAN1.Extracellular ZSCAN1 is captured by ZSCAN1-reactive B cells, which efficiently process and present cognate peptides to ZSCAN1-reactive CD4 T cells. Reciprocal antigen-specific B–CD4 T cell interactions sustain activation of both populations through peptide–MHC engagement, costimulatory signals, and cytokine exchange. This upstream circuit supports clonal persistence and expansion, plasma cell differentiation, and continued ZSCAN1 autoantibody production, while maintaining inflammatory conditions that support ongoing cytotoxic effector activity. ZSCAN1 autoantibody deposition, complement binding, and continued CD8 T cell activation amplify neuronal injury, generating a feed-forward loop in which additional tissue damage releases more antigen and reinforces the pathogenic B–CD4 T cell circuit. The model integrates the major observations in this study, including leptomeningeal immune expansion, intrathecal persistence of ZSCAN1-targeting B and CD4 T cell clones, and sustained ZSCAN1 autoantibody production and deposition within the brainstem.

Several observations support this framework. ZSCAN1-reactive B cell and CD4 T cell clones were strikingly enriched in the CNS and accounted for nearly all persistent and/or expanding antigen-linked B and CD4 T cell populations identified in CSF. Their convergence on shared regions of ZSCAN1 argues against these findings being a nonspecific byproduct of inflammation and instead points to a focused antigen-specific circuit operating within the diseased compartment. Even though few B cells were recovered from CSF, these included expanded, class-switched lineages encoding high-affinity ZSCAN1-binding receptors that persisted across time points despite intensive immunotherapy, suggesting that rare intrathecal B cells can nonetheless mark a durable and biologically important reservoir of disease-associated immunity.

Our tissue-based analyses argues that the relevant humoral response is not simply a diluted reflection of serum autoreactivity. IgG eluted from diseased brain was markedly restricted and regionally structured, with the brainstem repertoire dominated by a hypermutated lineage traceable to an expanded intrathecal clonotype. This anchors the antibody response directly within affected tissue rather than relying only on serum or recombinant inference, and it suggests that disease expression may reflect not only antigen specificity but also regional vulnerability. The hypothalamus and brainstem occupy specialized neurovascular and CSF-facing interfaces that support neuroendocrine exchange and autonomic regulation, potentially lowering the threshold for local immune access or effector injury. In such a setting, even a relatively sparse antigen-specific immune response could disproportionately disrupt homeostatic circuits controlling respiration, autonomic tone and endocrine function. Although antibodies to intracellular antigens are unlikely to initiate disease directly, our data suggest that once tissue injury has exposed antigen and enabled local antibody deposition, ZSCAN1-specific IgG may further exacerbate pathology. The antigen-specific co-purification of C3 and C4 with ZSCAN1-reactive IgG is consistent with a contribution of complement-mediated inflammation or tissue injury within the diseased CNS.

Beyond the disease biology, this study also addresses an important technical bottleneck. A central problem in autoimmune antigen discovery is that patient-derived BCR repertoires are now straightforward to recover, but experimentally testing large numbers of candidate antibodies remains slow and expensive. PAIRIS provides a practical solution to this scalability gap by using structure-based prediction to prioritize candidate antibody–antigen interactions before monoclonal synthesis and validation. In this study, PAIRIS successfully enriched for true ZSCAN1-reactive intrathecal clones and helped localize likely binding regions that were subsequently confirmed experimentally. Although computational protein folding prediction cannot yet replace biochemical validation, these results show that it can substantially narrow the search space and accelerate the path from repertoire sequencing to mechanistic insight.

More broadly, AEGIS provides an end-to-end framework for resolving autoimmune specificity from compartment to mechanism. Starting from CSF- and tissue-derived antibodies, it links high-resolution epitope mapping to patient-specific BCR and TCR clonotypes and anchors these findings within the diseased organ itself. Applied here, AEGIS defines ROHHAD as an antigen-driven neuroimmune disorder organized around a convergent ZSCAN1-specific B cell–CD4 T cell circuit in the CNS. More generally, this strategy should be applicable across a wide range of autoimmune and paraneoplastic diseases in which the relevant antigens, epitopes and pathogenic clones remain unknown, providing a generalizable blueprint for deconvoluting pathogenic immune circuits in human autoimmunity.

## Materials and Methods

### Patient Enrollment

Three out of the four patients described in the manuscript were enrolled a UCSF IRB-approved research study (13-12236). The final patient had remnant clinical samples sent to UCSF on a de-identified basis, which the UCSF IRB has determined does not require IRB review.

### PhIP-Seq

All PhIP-Seq was performed similar to our previously published multichannel protocol: https://www.protocols.io/view/derisi-lab-phage-immunoprecipitation-sequencing-ph-czw7x7hn?step=14.1

As previously described, the human peptidome library is a custom T7 phage display collection comprising 731,724 unique bacteriophage clones, each encoding a distinct 49–amino acid peptide displayed on the phage surface. Together, these peptides densely tile the annotated human proteome (including all isoforms available in 2016) using 25–amino acid overlaps. For each assay, 1 mL of the phage library was incubated with 1 µL of human serum overnight at 4 °C, followed by immunoprecipitation with 25 µL of a 1:1 mixture of protein A and protein G magnetic beads (Thermo Fisher, Waltham, MA; #10008D and #10009D). Beads were washed, and bound phage–antibody complexes were eluted into 1 mL of *E. coli* (BLT5403, EMD Millipore, Burlington, MA) at OD600 0.5–0.7 and amplified by incubation at 37 °C. The amplified phage pool was then subjected to a second round of incubation with the same individual’s serum, and the immunoprecipitation workflow was repeated. Phage DNA was extracted from the final enriched pool, barcoded, PCR-amplified, and ligated with Illumina adapters. Libraries were sequenced on an Illumina platform (Illumina, San Diego, CA) to a depth of ∼1 million reads per sample.

### Phage Assisted Scanning Epitope Recovery (PhASER)

A PhASER library was designed to interrogate the ZSCAN1 epitopes identified in the initial ROHHAD PhIP-Seq experiments. The library includes the 49-amino acid ZSCAN1 peptides found to be enriched in ROHHAD, and for each parent peptide we generated three complementary variant series. First, a single-residue scanning library was constructed in which each position was individually substituted with alanine, with native alanines replaced by glycine. Second, a truncation library was generated by introducing a stop codon at each position after residue 5, allowing progressive delineation of the minimal C-terminal sequence required for antibody recognition. Third, a linker-replacement library was generated in which increasing lengths of the peptide were progressively replaced with an XTEN sequence to delineate N-terminal boundaries, terminating 5 residues upstream of the C-terminus. To maximize sequence diversity across the library, the XTEN register was varied between parent peptides, and where XTEN replacement would otherwise preserve the native residue identity, that position was forcibly mutated to alanine or, for native alanines, glycine. The resulting oligonucleotide pool was synthesized and cloned into the phage display backbone as previously described. PhIP-Seq using the PhASER library was performed using the same protocol as for the human peptidome library.

### Antibody Tissue Extraction

1-5cm sections of midbrain, hypothalamus, and pons brain tissue were submerged in 0.5% BSA in D-PBS in 50mL conical tubes (Falcon) on ice. In a sterile polystyrene petri dish (Corning) the tissue was cut into 1-2mm fragments with a sterile scalpel on ice. The tissue was then gently mashed with the plunger of a sterile syringe on a 70uM cell strainer (Corning) over a 50mL conical tube. The strainer was washed with 1mL of 0.5% BSA in D-PBS. The antibody-containing flowthrough was collected and stored in a -80°C freezer.

### Sample Processing and Cryopreservation of PBMC and CSF cells

PBMCs were isolated from whole blood collected in CPT tubes (BD). Whole blood was centrifuged and PBMCs washed with D-PBS, then treated with 1x RBC lysis buffer (Biolegend, 420301). The PBMCs were washed twice with 1% BSA in D-PBS, stained with 0.04% Trypan Blue Solution (Gibco, 15250061) and counted using a hemocytometer. PBMCs were resuspended in 10% DMSO in FBS-HI. Fresh CSF was centrifuged in 15mL conical tubes (Falcon) and the supernatant was removed leaving a pellet of CSF cells. The CSF cell pellet was resuspended in RPMI-1640 with 40% FBS-HI with equal parts RPMI-1640 with 40% FBS-HI and 20% DMSO added dropwise. Both PBMCs and CSF cells were immediately placed in a - 80°C freezer in a freezing container (Mister Frosty, Thermo Fisher) and transferred to LN within 1-2 days.

### Single cell RNA sequencing

Cryopreserved, olig-conjugated antibody-stained CSF and PBMCs and fresh CSF cell suspensions were run on the 10x Genomics Chromium iX to form droplets and generate barcoded cDNA. Gene expression and BCR and TCR enriched V(D)J libraries were prepared according to the Chromium GEM-X Single Cell 5’ Reagent Kits v3(10x Genomics, CG000733) kit user guide. Library quality was observed using High Sensitivity D5000 ScreenTape Analysis (Agilent) and quantified using Lib Quant Kit (Illumina/Uni) (Roche, 07960140001). Sequencing was performed on the NovaSeq X (Illumina) with NovaSeq™ X Series 25B Reagent Kit (100 Cyc)100-cycle Kit (Illumina, 20125967). Libraries were sequenced at a target depth of 20,000 read pairs/cell for 5’ Gene Expression libraries and 5,000 read pairs/cell for V(D)J libraries. Demultiplexed FastQ files were uploaded and aligned in 10x Cell Ranger 9.0.1.

### AIM Assay

The AIM assay was performed as previously described(*23*, *38*). In brief, patient PBMCs were thawed, washed, and equilibrated in RPMI with 10% FBS for 2 hours at 37°C. PBMCs were washed and resuspended in serum-free RPMI and plated at 1 x 10^6^ cells per well in 96-well round-bottom plates. PBMCs were stimulated with a ZSCAN1 peptide pool (107 15mer peptides overlapping by 11 amino acids, JPT Peptide Technologies) at a final concentration of 1 µg/ml per peptide or 0.2% DMSO vehicle control as a negative control. As positive control, PBMCs were also stimulated with 2 µg/ml phytohemagglutinin-L. PBMCs were stimulated for 24 hours, washed with FACS wash buffer (1X DPBS without calcium or magnesium, 0.1% sodium azide, 2 mM EDTA, 1% FBS) and stained with the following antibody panels: CD3 Alexa 647 (clone OKT3, 317312, BioLegend), CD4 Alexa 488 (clone OKT4, 317420, BioLegend), CD8 Alexa 700 (clone SK1, 344724, BioLegend), OX-40/CD134 PE-Dazzle 594 (clone ACT35, 350020, BioLegend), CD69 PE (clone FN-50, 310906, BioLegend), CD137/4-1BB BV421 (clone 4B4-1, 309820, BioLegend), CD14 PerCP-Cy5 (clone HCD14, 325622, BioLegend), CD16 PerCP-Cy5 (clone B73.1, 360712, BioLegend), CD19 PerCP-Cy5 (clone HIB19, 302230, BioLegend) and Live/Dead Dye eFluor 506 (65-0866-14, Invitrogen). Cells were washed with FACS wash buffer, fixed with 2% paraformaldehyde (BD), and stored in FACS wash buffer in the dark at 4°C until ready for acquisition on an LSR Fortessa (BD) flow cytometry. Flow cytometry data were analyzed with FlowJo (BD). The gating strategy is shown in Fig. S11. The frequencies of ZSCAN-specific T cells were calculated by subtracting the no stimulation background from the corresponding ZSCAN pool-stimulated samples.

### Tissue staining and TSA amplification

Tissue sections were prepared using standard methods, rehydrated in phosphate-buffered saline (PBS), and processed as needed for antigen retrieval. Endogenous peroxidase activity was quenched by incubating sections in 0.3% H₂O₂ in PBS for 10 min at room temperature, followed by three washes in PBS. Sections were then blocked in PBS containing 5% BSA and/or 5% Donkey serum with 0.3% Triton X-100 for 60 min at room temperature. Endogenous biotin was blocked using a commercial biotin-blocking kit according to the manufacturer’s instructions (E-21390, ThermoFisher scientific).

Sections were incubated with biotin-conjugated anti-human IgG diluted in blocking buffer overnight at 4°C. The next day, sections were washed three times in PBS and incubated with streptavidin–horseradish peroxidase (HRP) for 20–30 min at room temperature. Sections were washed three times in PBS and developed using a TSA Alexa Fluor 594 working solution prepared immediately before use. TSA development was performed for 5 min at room temperature, and the reaction was stopped by washing extensively in PBS. Sections were mounted in antifade mounting medium with DAPI and imaged on a laser-scanning confocal microscope. Alexa Fluor 594 signal was collected in the 594 channel using identical acquisition settings across samples, and DAPI was collected in the blue channel with appropriate excitation/emission settings.

### Antibody immunoprecipitation

Finely mapped peptides derived from PhASER or PhIP-Seq fragments were synthesized at >95% purity with an N-terminal biotin-Ahx modification and GGSG linker (Biotin-Ahx-GGSG-peptide; Lifetein Inc.). Biotinylated peptides were added sequentially to either 1 mL of cerebrospinal fluid (CSF) in 1× sample buffer (SB) or 0.2 mL of serum in 1× SB and incubated with end-over-end rotation for 30 min at room temperature.

Dynabeads M-280 Streptavidin (Thermo Fisher Scientific) were washed with 1× PBS and resuspended to the original volume. Washed beads (100 µL) were added to each sample and incubated with end-over-end rotation for 30 min at room temperature. Samples were placed on a magnetic rack, and unbound material was transferred to a new tube. Beads were washed with two volumes of affinity purification buffer (50 mM Tris-HCl, pH 7.4; 150 mM NaCl; 1 mM EDTA; 0.5% Igepal NP-40), then frozen at −80 °C for downstream mass spectrometry analysis.

Due to limited patient material, peptide affinity purifications were performed sequentially in the following order: ZSCAN1 fragment 15, fragment 2, fragment 13, and fragment 17. For capture of the total antibody repertoire, Protein A/G Dynabeads were incubated with 1 mL of CSF or brain tissue supernatant, then washed and frozen using the same procedure described above.

### Monoclonal antibody generation

Recombinant monoclonal antibodies were generated by GeneScript (Piscataway, NJ). Variable region sequences of the immunoglobulin heavy and light chains were synthesized and cloned into human IgG1 expression vectors containing the appropriate constant regions and light chain isotype (kappa or lambda) was selected according to the BCR sequencing data. Antibodies were transiently expressed in the GeneScript TurboCHO™ expression system according to the manufacturer’s protocols. Following expression, antibodies were harvested from culture supernatants and purified antibodies were buffer-exchanged into sodium acetate buffer, and 0.2M L-Arginine, pH 5.5 and assessed for purity by CD-SDS and SEC-HPLC.

### Antibody repertoire mass spectrometry (Ig-MS)

Affinity-captured proteins were prepared either by protein A/G immunoprecipitation or by peptide-affinity capture resin and subjected to peptide-sequencing by mass spectrometry. For protein A/G immunoprecipitated samples, captured antibody Fab domains and potential co-bound antigen protein candidates were released by pre-treatment with IdeS (Sino Biological, Cat# 40610-S07E, 40 units, 37 deg C, 1h) under native reaction conditions. The released material as well as the bead bound remnant samples were then denatured (4M urea), reduced (10 mM DTT, 30 min), and alkylated (15 mM iodoacetamide, 60 min), followed by 5 mM DTT quench, and then subjected to further digestion with sequencing grade proteases as indicated in each sample. Fab-containing samples were divided into equal-volume reactions and diluted to 1M urea with appropriate reaction buffers for digestion with trypsin (Promega, V5280), chymotrypsin (Promega, V1061), or Krakatoa (Cinder Biological, Inc., CB23726) following manufacturers’ specifications. The remnant bead-bound material, was digested with trypsin.

The resulting peptide samples were desalted with C18 zip tips, and then analyzed by tandem liquid chromatography mass spectrometry (LC-MS/MS) on a QExactive mass spectrometer (Thermo), equipped with a 10,000-psi system nanoACUITY (Waters) ultra-high performance liquid chromatography (UPLC) instrument for reversed-phase C_18_ chromatography. Mass lists were converted with in house software called PAVA, and analyzed using ProteinProspector v. 6.8.1(*43*). Database searches were performed with 20 ppm precursor, 30 ppm fragment mass error tolerances, carbamidomethylation (Cys) as fixed modification, and as variable modifications: oxidation (Met), deamidation (Gln, Asn), acetylation (N-term), N-term Met loss, pyroglutamate (Gln), carbamylation (Cys, Lys). Data were searched against the SwissProt human database (downloaded 2024.01.24) containing 42,506 entries, concatenated with a fully randomized set of protein entries, for estimation of false discovery rate; FDR was set to 1%(*44*). Data were also searched against a patient repertoire of immunoglobulin sequences in a library of 430 entries using score thresholds established from SwissProt searches, with maxiumum E-values for protein (0.01) and peptide (0.001); all matched MS/MS spectra, particularly from CDR3 regions, were manually validated. Unique peptide sequences are defined here as matching uniquely to individual entries in the patient repertoire library, and those peptides that passed CDR3-identity filtering were used to infer enrichment. Normalized spectral counting was used as an approximation of relative peptide and protein abundances.

### Immunoprecipitation mass spectrometry (IP-MS)

To identify co-immunoprecipitated proteins by patient antibodies, immunocomplexes from patient tissues and biofluids were captured with magnetic protein A/G beads, followed by gentle washes of 0.1% NP40 in PBS buffer, then subjected to proteomic analysis. Samples were denatured, reduced and alkylated as above, then divided into 2-3 equal volume reactions and digested on bead with proteomics grade proteases including trypsin, chymotrypsin, and AspN or Krakatoa, and fully analyzed as described above.

### Biolayer interferometry (BLI)

Binding kinetics and dissociation constants (Kd) were measured using biolayer interferometry on the Gator Prime system. Biotinylated peptides were immobilized on streptavidin-coated biosensors and incubated with serial dilutions of monoclonal antibody analytes. Initial scouting experiments were performed to determine optimal peptide loading densities on the biosensors to ensure measurements within the kinetic range. All experiments were conducted in Gator Kinetics Buffer. Biosensors were equilibrated by incubation in buffer for 10 min, followed by a 30 s pre-wetting step with shaking at 1000 rpm. The assay sequence consisted of a 120s baseline, 120s peptide loading, a second 120s baseline, a 120s association phase with monoclonal antibody, and a 600s dissociation phase in buffer. Kinetic parameters were determined by globally fitting the association and dissociation data using a 1:1 binding model, from which dissociation constants (Kd) were calculated. Peptide sequences used for monoclonal antibody binding measurements are listed in the table below. Sequences include an N-terminal GGSG linker.

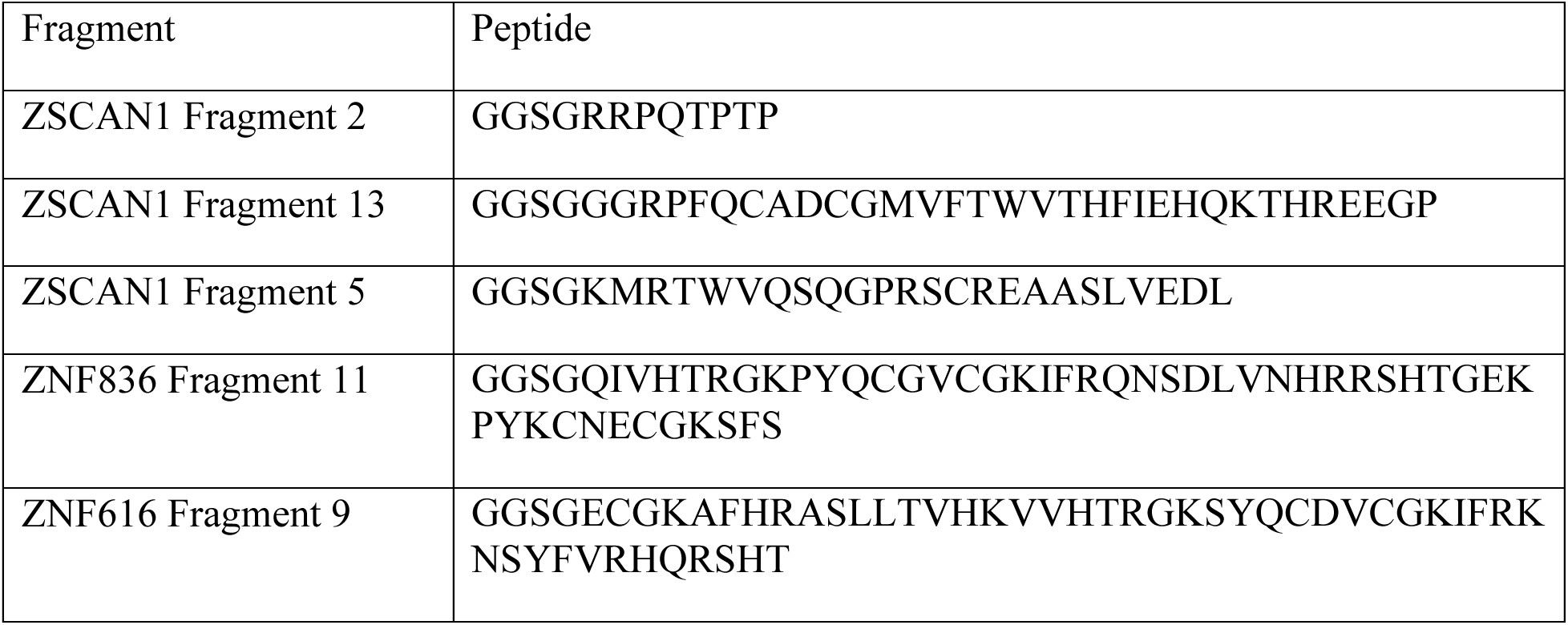

### Prediction of Antibody-antigen Interaction at high Resolution In Silico (PAIRIS)

PAIRIS is a computational framework for predicting antibody-antigen binding specificity that integrates structure prediction and binding energy estimation. The pipeline takes as input a peptide sequence and the heavy and light chain variable domains of a monoclonal antibody, which are provided to AlphaFold3 (AF3) for structure prediction. Binding confidence is assessed using the interface predicted template modeling (iPTM) score, calculated as the average iPTM between the peptide and each antibody chain. Predicted structures are then evaluated using Rosetta to compute estimated binding free energies (ΔG).

### Structure prediction and confidence scoring

AlphaFold3 (AF3) was used to predict structures of antibody:peptide complexes for all patient-derived IgG BCR clonotypes against tiled ZSCAN1 peptides. Input consisted of three protein chains: the variable domains of the heavy and light chains of a given BCR and a k-length peptide (k=15 or k=25) from the 408-amino acid ZSCAN1 protein (UniProt: Q8NBB4-1). Peptides were generated by sliding-window tiling in 1-amino acid increments, yielding 394 15-mer and 382 25-mer peptides. For the initial screen of 81 IgG BCR clonotypes, 25 model seeds were used per prediction (5× the default), as increased seed number significantly improves accuracy for antibody-antigen complex predictions(*33*). Additional AF3 predictions were generated for mAb2 complexes with 15-mer peptides from ZNF616 (UniProt: Q08AN1, aa 217-264) and ZNF836 (UniProt: Q6ZNA1, aa 265-312) using 100 model seeds.

### Binding energy calculations

Rosetta binding energies (ΔG) were computed for AF3-predicted antibody:peptide structures using PyRosetta’s InterfaceAnalyzerMover with the ref2015 scoring function(*35*, *45*, *46*). The peptide (chain A) was designated as one partner and the antibody heavy and light chains (chains B and C) as the other partner (docking partners string “A_BC”). To account for side-chain optimization, both the bound complex and separated partners were repacked prior to energy evaluation (pack_input=True, pack_separated=True), allowing side chains to adopt favorable rotamers in both bound and unbound states. Final ΔG values were reported in Rosetta Energy Units (REU), with more negative values indicating stronger predicted binding interactions.

### Prioritization of candidate BCRs

BCRs predicted to bind ZSCAN1 were identified using thresholds iPTM > 0.85 and negative Rosetta ΔG.

### *In silico* deep mutational scanning

To assess the tolerance of mAb2 binding to sequence variation within its ZSCAN1 epitope, computational deep mutational scanning was performed using AF3. The wild-type peptide sequence GRPFQCADCGMVFTWVTHFIEHQKTHREE was used as the reference peptide. All possible single-amino acid substitutions were generated at each position in the peptide, yielding 551 variant peptides (19 alternative amino acids × 29 positions). AF3 predictions were generated for mAb2 in complex with the wild-type peptide and each variant using 25 model seeds per complex. Changes in iPTM score relative to wild-type were used to evaluate the effect of each substitution on predicted binding affinity.

### TCR reconstruction and expression

Paired TCRα and TCRβ sequences from expanded CSF CD4 T cell clonotypes were reconstructed from single-cell sequencing data and expressed in a Jurkat-based reporter system. TCR α/β sequences were encoded in a single expression cassette separated by a P2A self-cleaving peptide and introduced by lentiviral transduction. Recipient cells were TCRα/β-deficient Jurkat cells (TRAC and TRBC1 knockout) engineered to express human CD4, as previously described(*39*). Stable TCR-expressing lines were generated and expanded prior to functional assays.

### HLA expression and antigen-presenting cells

Patient-derived HLA class II alleles were synthesized based on IMGT reference sequences and individual allele pairs were cloned into lentiviral expression vectors in α–β orientation under the control of the Pgk promoter, with a P2A linker and IRES-Neo selectable marker to enable coordinated expression and stable selection. Constructs were introduced into K562 cells expressing the CIITA transcription factor and lacking endogenous HLA-II expression to generate engineered antigen-presenting cells expressing individual HLA class II allele pairs.

### Antigen presentation and reporter assay

Antigen-specific activation was assessed using an activation-dependent reporter system based on NFAT-driven mCherry expression(*47*). TCR-engineered Jurkat reporter cells were co-cultured with HLA-expressing K562 antigen-presenting cells in the presence of ZSCAN1-derived peptides fused to human CD74 to facilitate class II presentation(*40*). Reporter activation was quantified by flow cytometry as a measure of TCR engagement: mCherry-positive reporter cell population was calculated as a percent of the total reporter cell population.

### Epitope mapping and validation

Candidate regions were initially evaluated using overlapping 24–amino acid ZSCAN1-derived peptides. Minimal activating sequences were subsequently defined using shorter 15–16 amino acid peptides in the context of the relevant HLA class II allele pairs. TCR activation across peptide–HLA combinations enabled assignment of cognate peptide–MHC ligands and refinement of minimal epitopes for individual clonotypes.

### Analysis

#### PhIP-Seq

As previously described(*48*), next generation sequencing reads from fastq files were aligned at the level of amino acids using RAPSearch2(*49*). All human peptidome analysis was performed at the peptide level. Reads were normalized to the read depth for each individual sample by calculating RPK (reads per 100k). For the PhIP-Seq of the high-resolution epitope mapping (PhASER), the RPK for each peptide was compared to the RPK of each peptide in the input library, to calculate the enrichment above input library.

#### Single-cell RNA sequencing

Raw scRNA-seq, scTCR-seq, scBCR-seq, and cell-surface protein (CSP) data were processed with Cell Ranger (v9.0.0; 10x Genomics) using cellranger multi(*50*). scRNA-seq and V(D)J data were aligned to GRCh38-2024-A and GRCh38-alts-ensembl-7.1.0 reference packages. CSP (TotalSeq-C) counts were processed using a custom feature reference containing the surface-marker panel. Downstream integration and analysis were performed in R (v4.5.0) using Seurat (v5.2.1), Azimuth (v0.5.0), and DoubletFinder (v2.0.4)(*51–53*). V(D)J contigs from Cell Ranger were further analyzed with the Immcantation framework (v4.5.0). Clonal families were assigned with Change-O (v1.3.0), using amino-acid–based distance models: a distance threshold of 0 for TCRs and 0.15 for BCRs(*54*). Gene expression matrices were combined into a single Seurat object. Cells with >400 detected transcripts were retained. Counts were normalized per sample with sctransform (v0.4.2) and integrated using shared anchors; TCR V genes were excluded from normalization and clustering. Immcantation-derived BCR/TCR annotations were linked to the scRNA-seq object via shared cell barcodes to generate paired expression–repertoire objects, which were then subsetted for B- and T-cell–focused analyses.

#### BCR analysis

Cell Ranger all contig fasta and associated annotation files from 10x Genomics BCR libraries were uploaded to the PipeBio platform and processed using the 10x Genomics Analysis workflow. The workflow consists of running 10x QC with a UMI count cutoff of 3 and read count cutoff of 50 reads, and IMGT v1.1 germline annotations. Clonotypes were defined based on >85% identity heavy-chain CDR3 amino acid sequences and shared V and J gene usage. Paired light chains were retained for downstream characterization. For CSF B cells, 10x gene expression profiles were linked to paired BCR sequences to obtain joint expression and BCR information at the single-cell level. This enabled identification of ZSCAN1-reactive monoclonal antibodies (mAbs) with their corresponding B-cell transcriptional states. Linked gene expression data were available for clone 11 and clone 12, and for 3 of 6 cells in clone 2. CSF B cells with paired expression data were annotated using Azimuth with the default human B-cell reference to infer differentiation states. ZSCAN1-positive cells were visualized in a heatmap showing Azimuth-predicted differentiation state alongside normalized expression of canonical marker genes for each state.

#### B cell V sequence similarity tree construction

Heavy-chain sequences were processed with the Immcantation pipeline starting from Cell Ranger V(D)J outputs and FASTA files. Sequences were annotated with IgBLAST, clonally assigned using a distance threshold of 0.15, and germlines inferred with CreateGermlines(*55*). V-gene regions were extracted using annotated germline coordinates, producing a table of sequence IDs and V-gene germline sequences. Pairwise distances between V-gene germline sequences were computed using the TN93 substitution model with pairwise deletion to accommodate minor length variation. Distance trees were visualized with ggtree in R.

#### Solvent-accessible surface area and minimum-distance calculations for mAb2

To evaluate the AlphaFold-predicted interaction between mAb2 and ZSCAN1 fragment 13, per-residue changes in solvent-accessible surface area (ΔSASA) were computed for the ZSCAN1 fragment using PyMOL(*56*). The AlphaFold-predicted mAb-B–ZSCAN1 complex was used to calculate SASA in the bound state and, separately, for the unbound ZSCAN1 fragment. SASA was computed with a 1.4 Å solvent probe radius using the dot solvent representation (dot density 5). Per-atom ΔSASA was calculated as unbound minus bound SASA, then summed across atoms within each residue to yield per-residue ΔSASA. Residues with larger decreases in SASA upon binding were interpreted as candidate contact residues. In addition, minimum distances between ZSCAN1 fragment residues and the mAb-B heavy and light chains were computed from the CIF structure. Atom coordinates for the peptide and antibody chains were parsed (MMCIFParser), and for each peptide residue the minimum interatomic distance to the heavy chain and to the light chain was recorded.

#### T cell analysis

For CSF T cells, 10x gene expression profiles were linked to paired TCR sequences to obtain joint expression and TCR information at the single-cell level. This enabled extraction of transcriptional profiles for ZSCAN1-validated TCR clonotypes to infer T-cell subtype and helper-program signatures. An initial CD4/CD8 assignment was performed using CD8A and/or CD8B expression (CD8 if either gene was expressed; otherwise CD4), and was cross-checked by visualizing clonal expansions against CD4/CD8 marker expression. This yielded a set of 421 CSF CD4 T cells for downstream helper-program analysis.

Helper-program scoring was performed with AUCell(*57*). First, AUCell was run across the 421 CD4 cells using the default setting of the top 5% of expressed genes and the default global AUC threshold to assign a single best-matching CD4 subtype when multiple subtypes exceeded threshold. AUCell was then applied to canonical gene sets representing active helper programs and “primed” program signatures. For these more granular signatures, the top 10% of expressed genes were used, and multiple programs were permitted per cell because CD4 T cells can exhibit overlapping activation/priming states. Of nine persistent TCR clones, five were tested and four were ZSCAN1-validated. The 15 cells belonging to the four ZSCAN1-validated clones were used for final visualization of CD4 differentiation subtype and helper-program scores.

For differential gene expression (DGE), the Seurat object containing the 421 CD4 T cells was normalized, scaled, and subjected to PCA. UMAP was constructed using the first 15 principal components. Outliers were removed based on UMAP distance (threshold 10, chosen due to a clear separation beyond this value), yielding 349 cells and 10 remaining ZSCAN1-validated cells. DGE was performed using Seurat FindMarkers with a Wilcoxon rank-sum test (min.pct = 0.5, logfc.threshold = 0.25). Results were visualized by plotting, for each gene, the difference in the fraction of cells expressing the gene between groups against the average log2 fold-change.

## Acknowledgement

Sequencing was performed at the UCSF CAT, supported by UCSF PBBR, RRP IMIA, and NIH 1S10OD028511-01 grants.

## Data, code, and materials availability

All relevant data and code will be made publicly available upon publication.

## Extended Data

**Extended Data Fig. 1.**
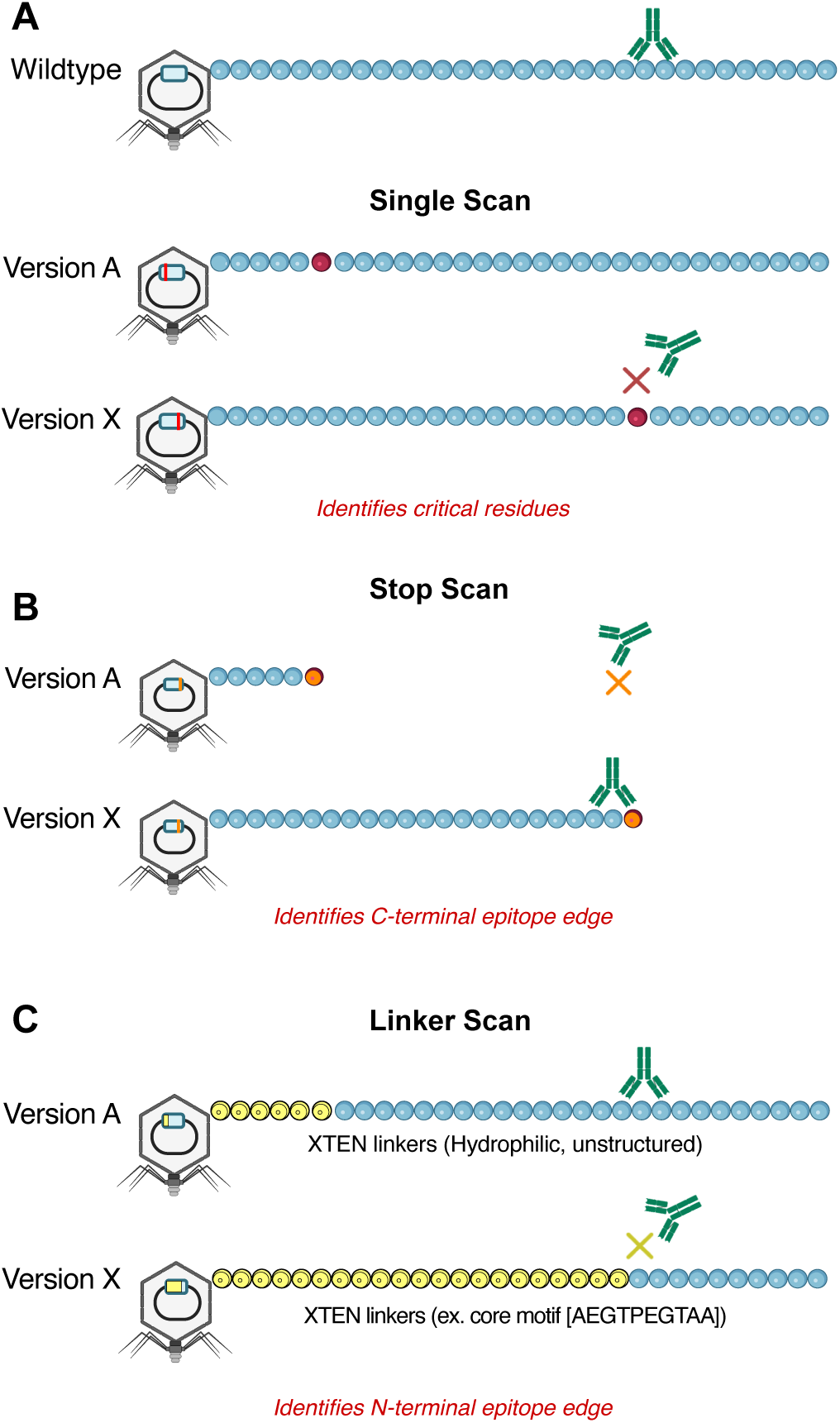
Phage-assisted scanning epitope recovery (PhASER) design A) Single–residue scan. For each PhIP-seq–positive 49-mer, every position is mutated individually to alanine (or to glycine when alanine is the native residue). Antibody binding to the wild-type fragment is compared with binding to each single–amino-acid variant; loss of binding identifies residues essential for epitope recognition. **B) Stop scan.** To define the C-terminal boundary, each codon is sequentially replaced with a stop codon to generate progressively truncated fragments. Binding is observed only when the displayed fragment fully contains the epitope, enabling precise localization of the C-terminal edge. **C) Linker scan.** To define the N-terminal boundary, an N-terminal XTEN linker is extended one residue at a time. Binding drops abruptly when the linker encroaches into the epitope, revealing the N-terminal edge. XTEN was selected for minimal immunogenicity and limited structural interference (hydrophilic, unstructured).

**Extended Data Fig. 2.**
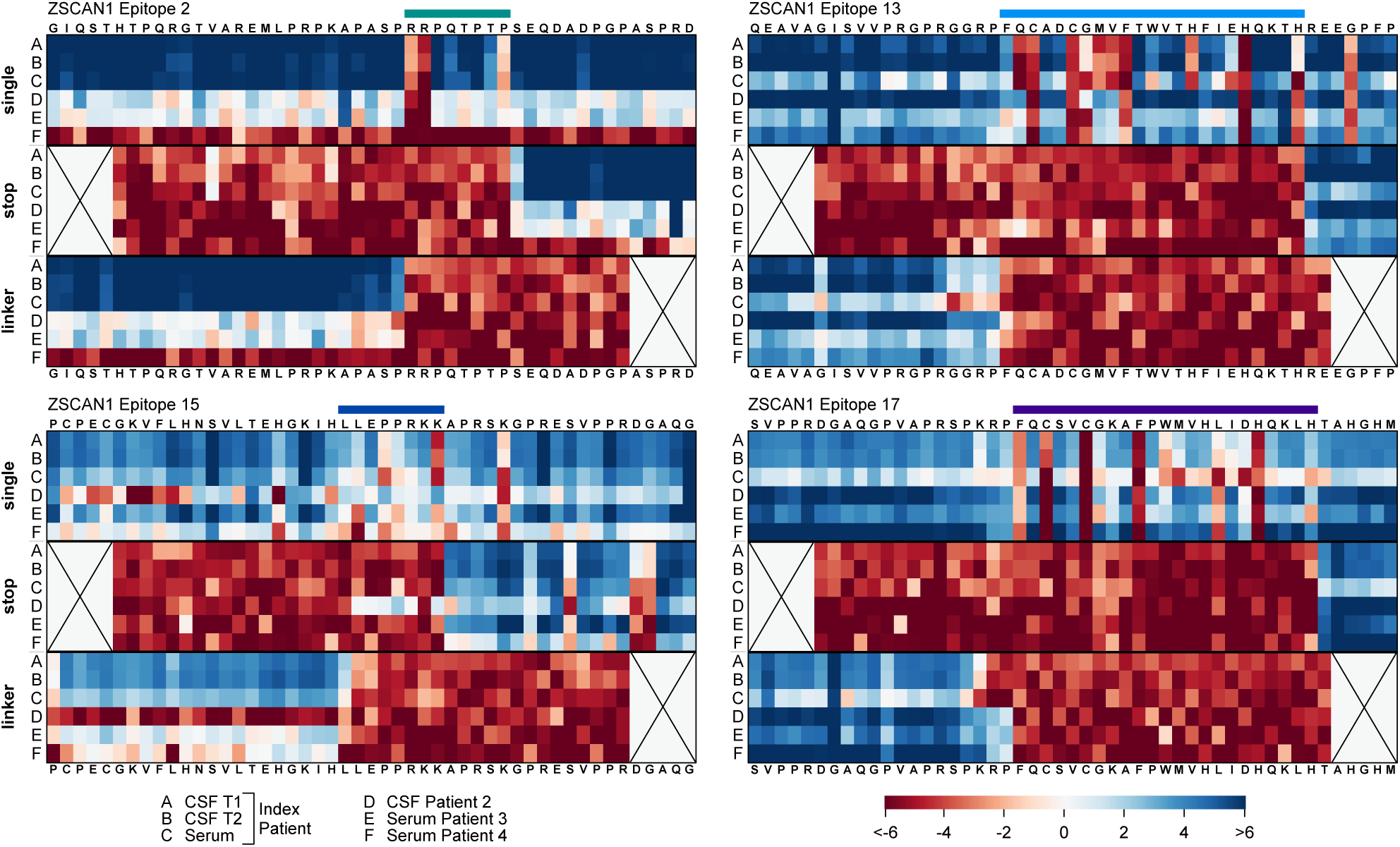
Complete PhASER maps of ZSCAN1 autoantibody epitopes. Complete PhASER epitope maps for the four most reactive ZSCAN1 peptides identified by PhIP-seq (Figure 2). Each cell in the ‘single’ scan shows results from mutating individual amino acid residues to alanine (or to glycine if alanine was the native residue). The ‘stop’ scan shows results from mutating each residue to a stop codon, starting from the sixth residue, to define the C-terminal limit of the epitope. The ‘linker’ scan shows results from progressive N-terminal XTEN linker extension, one residue at a time until the fifth-to-last residue, to define the N-terminal boundary of the epitope; each cell corresponds to the peptide being mutated to the XTEN linker up to and including the respective residue. Signal is calculated as log2 fold-change over the input library. Rows A,B, and C represent index patient samples from serum, CSF #1, and CSF #2, respectively; rows D, E, and F represent samples from three additional patients with ROHHAD. The coordinates of each ZSCAN1 peptide fragment containing the epitopes are as follows: epitope 2, aa 25-73; epitope 13, aa 313-361; epitope 15, aa 361-409; epitope 17, aa 400-448 (ZSCAN1 isoform X2, NCBI: XP_006723212.1). Amino acid coordinates of epitopes presented in Figure 2 come from the canonical ZSCAN1 sequence (UniProt: Q8NBB4).

**Extended Data Fig. 3.**
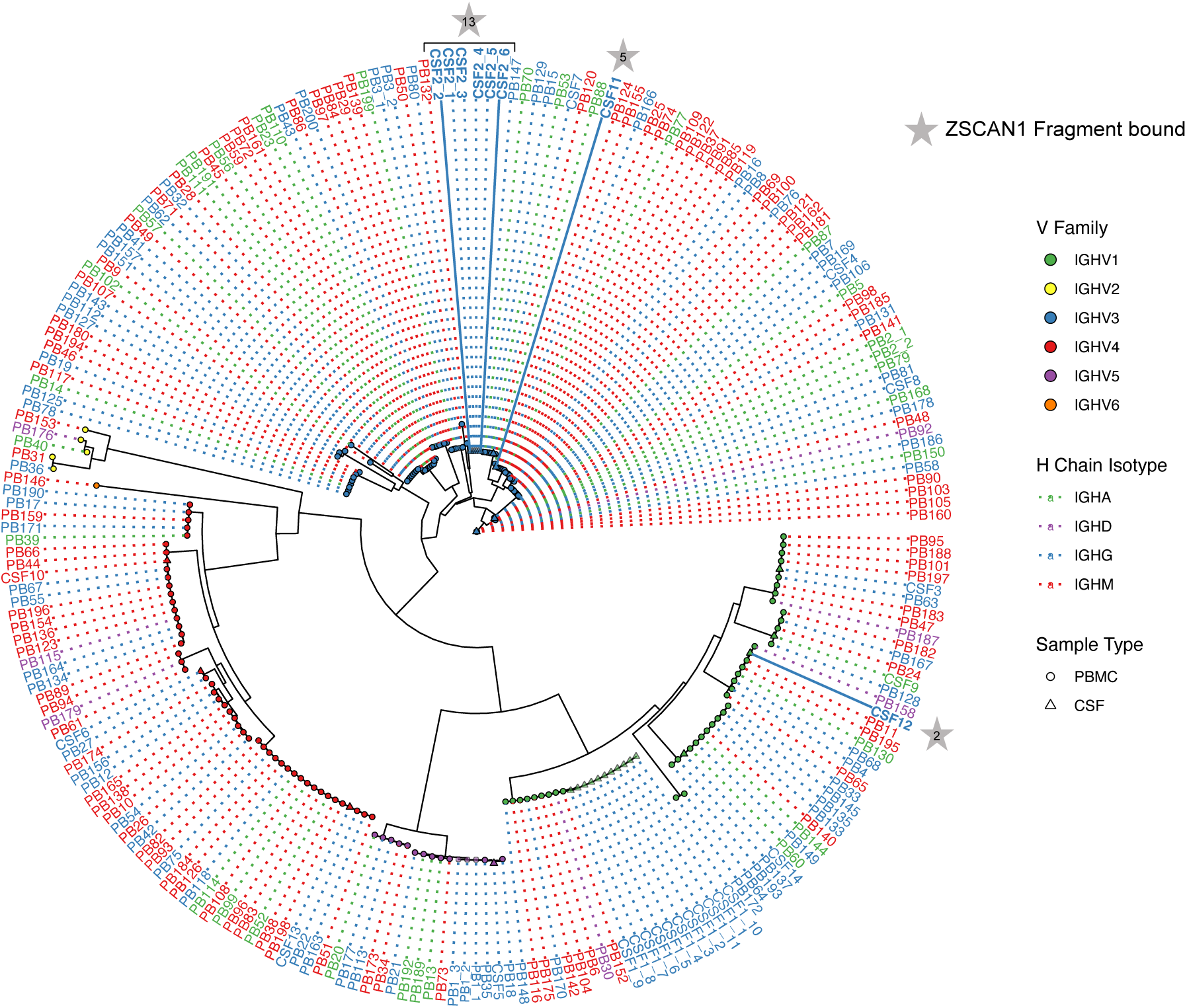
Heavy-chain V germline similarity tree. Distance tree of monoclonal antibody heavy-chain V gene germline sequences constructed using the TN93 substitution model with pairwise deletion. Tree nodes are colored by V-gene family, shaped by sample type (CSF vs PBMC). Branch annotations and dotted line colors denote heavy-chain isotype. BCR clones from the main text are labeled as follows: Clone A = CSF12, Clone B = CSF2, Clone C = CSF11, Clone D = CSF4, Clone E = CSF5, Clone F = CSF9, Clone G = CSF13.

**Extended Data Fig. 4.**
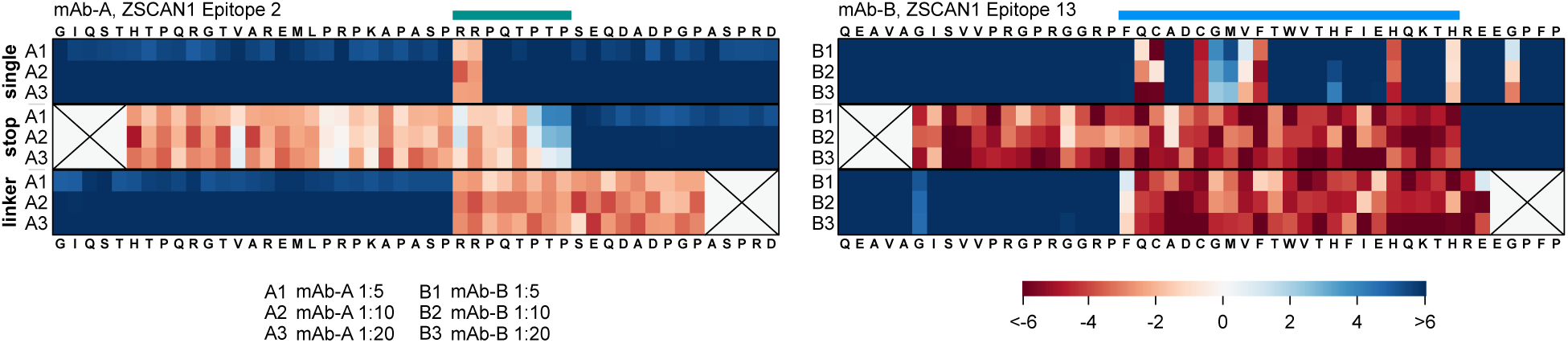
Complete ZSCAN1 epitope PhASER maps of mAb-A and mAb-B. Complete PhASER epitope maps for patient-derived monoclonal antibodies mAb-A (ZSCAN1 Epitope A, left) and mAb-B (ZSCAN1 Epitope B, right). Each cell in the ‘single’ scan shows results from mutating individual amino acid residues to alanine (or to glycine if alanine was the native residue). The ‘stop’ scan shows results from mutating each residue to a stop codon, starting from the sixth residue, to define the C-terminal limit of the epitope. The ‘linker’ scan shows results from progressive N-terminal XTEN linker extension, one residue at a time until the fifth-to-last residue, to define the N-terminal boundary of the epitope; each cell corresponds to the peptide being mutated to the XTEN linker up to and including the respective residue. Signal is calculated as log2 fold-change over the input library. Rows represent different dilutions of the monoclonal antibodies (1:5, 1:10, 1:20). The coordinates of each ZSCAN1 peptide fragment are consistent with Extended Data Fig. 2.

**Extended Data Fig. 5.**
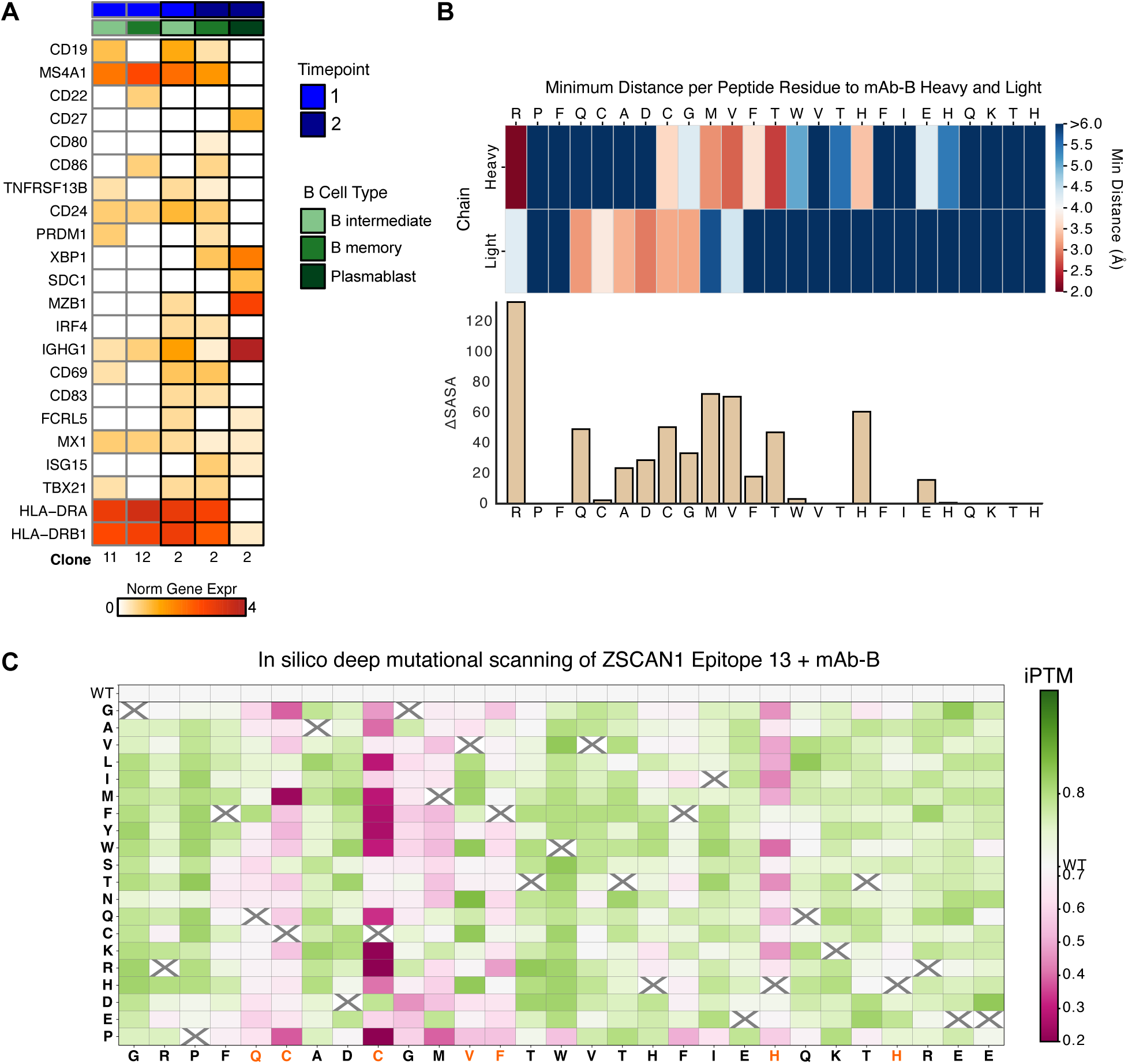
ZSCAN1 reactive B cell clone evaluation. **A)** Azimuth-based prediction of B-cell differentiation states for ZSCAN1 positive clones, with a heatmap showing normalized expression of marker gene sets across timepoints for intermediate, memory, and plasmablast states. Intermediate indicates gene expression profiles spanning both naive and memory states. **B)** Per-residue delta SASA (solvent-accessible surface area), measuring loss of surface area upon binding, of ZSCAN1 Epitope 13 in complex with mAb-B as predicted by PAIRIS is shown alongside a heatmap of the minimum distances to heavy and light chains, highlighting residues most engaged with each chain. **C)** In silico deep mutational scanning of ZSCAN1 Epitope 13 in complex with mAb-B, showing iPTM scores for each residue substitution as predicted by PAIRIS. Critical epitope residues as determined by PhASER are highlighted in orange.

**Extended Data Fig. 6.**
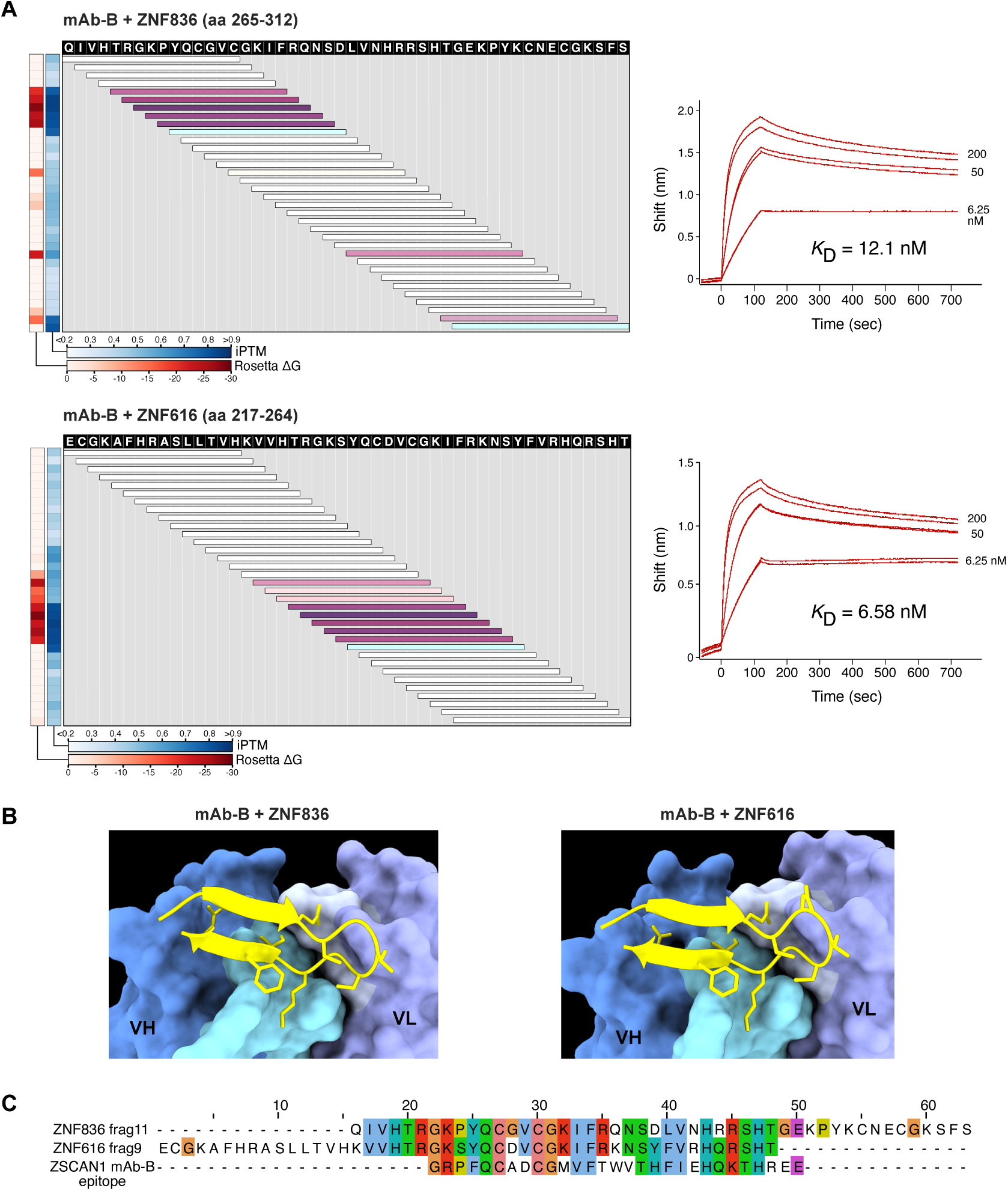
Binding characteristics of mAb-B. **A)** Validation of PAIRIS binding predictions by PhASER and biolayer interferometry (BLI) for mAb-B against ZNF836/ZNF616 peptides. Heatmaps display iPTM scores (average interface confidence between peptide and both antibody chains) and Rosetta ΔG values for 15-mer peptides spanning ZNF836 aa 265-312 (top, UniProt: Q6ZNA1) and ZNF616 aa 217-264 (bottom, UniProt: Q08AN1). Horizontal bars represent overlapping peptides spanning the indicated amino acid positions, with the color representing the sum of iPTM and Rosetta values; dashed lines denote PhASER epitope regions. BLI sensorgrams using displayed 48AA peptides confirm binding with calculated KD values. **B)** AlphaFold3-predicted structures show the highest-scoring peptide-antibody complex, with sidechain atoms within 4Å displayed. **C)** Multiple sequence alignment of ZNF836 and ZNF616 peptides with the ZSCAN1 epitope for mAb-B.

**Extended Data Fig. 7.**
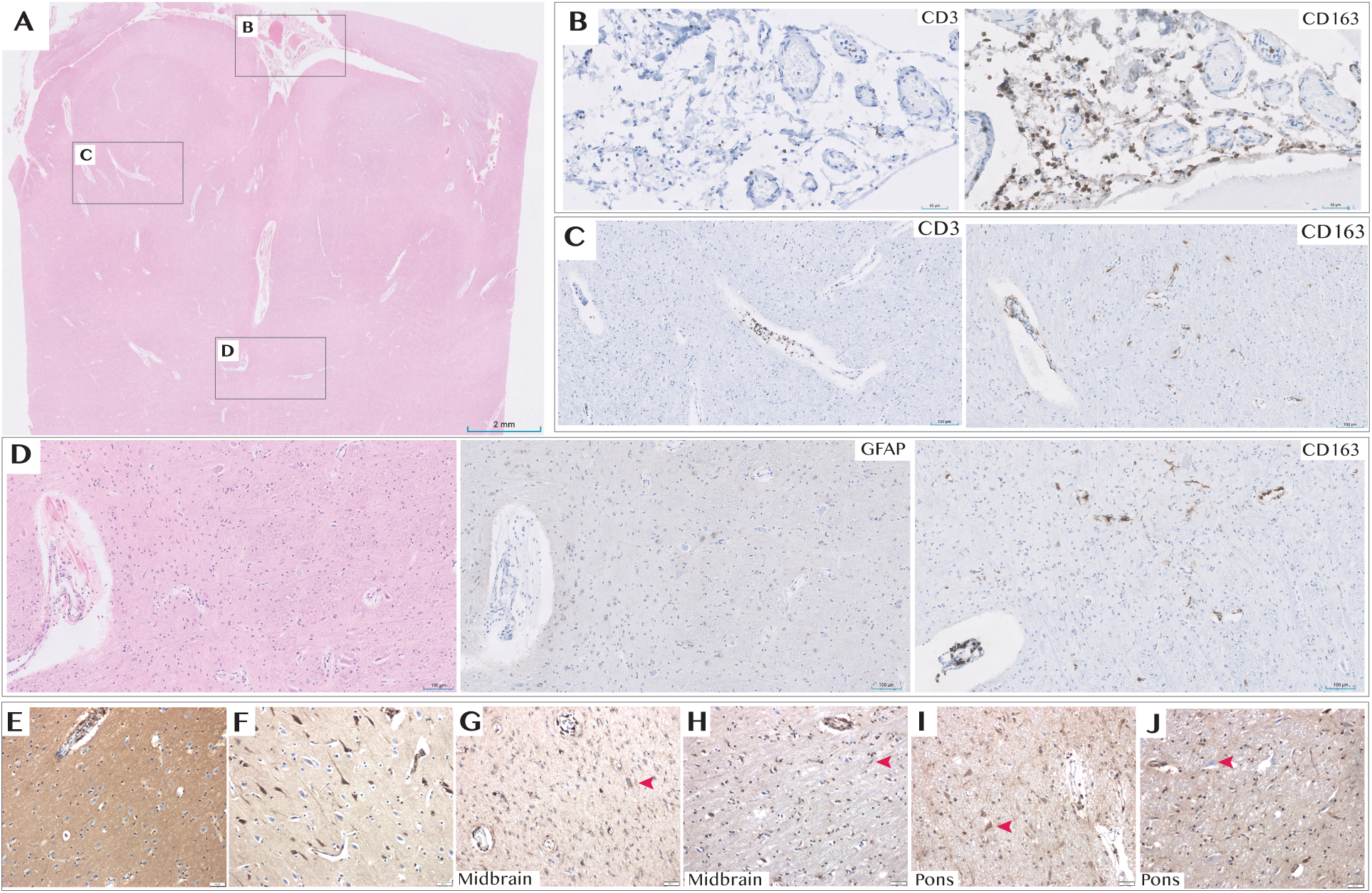
Key Neuropathology Findings. Autopsy showed widespread acute and chronic hypoxic–ischemic encephalopathy (most consistent with recurrent respiratory arrest) with thickened, opaque leptomeninges. Microscopy revealed expansion of leptomeningeal and perivascular (Virchow–Robin) spaces by a sparse inflammatory infiltrate composed predominantly of macrophages and T cells, seen across cortex, subcortical and periventricular white matter, hypothalamus, basal ganglia and brainstem; CD20 and CD138 immunohistochemistry was essentially negative (data not shown). **A)** low-power H&E overview with boxed regions shown in **B–D**. **B)** leptomeningeal inflammatory infiltrate with CD3-positive T cells (left) and CD163-positive macrophages (right). **C)** dilated perivascular spaces containing CD3-positive T cells (left) and CD163-positive macrophages (right), with scattered CD163-positive reactive microglia in the parenchyma. **D)** region with acute neuronal necrosis showing patchy reactive astrogliosis (GFAP, middle) and microglial activation/microgliosis (CD163, right) adjacent to the H&E section (left). **E–J)** IgG immunohistochemistry (clone RWP49, Leica; human IgG1; 15-min incubation on BOND-PRIME): control cortex shows weak-to-absent neuronal cytoplasmic IgG (E), whereas ischemic neurons in the same control tissue show variable weak-to-strong staining (F). Patient midbrain (G) and pons (I) show increased neuronal cytoplasmic IgG staining compared with location-matched controls (H, control midbrain; J, control pons); red arrowheads indicate representative neurons. Control tissue in **H** and **J** was obtained in the setting of acute bacterial meningitis with mixed acute and chronic inflammation involving leptomeninges and perivascular spaces. Scale bars, as indicated.

**Extended Data Fig. 8.**
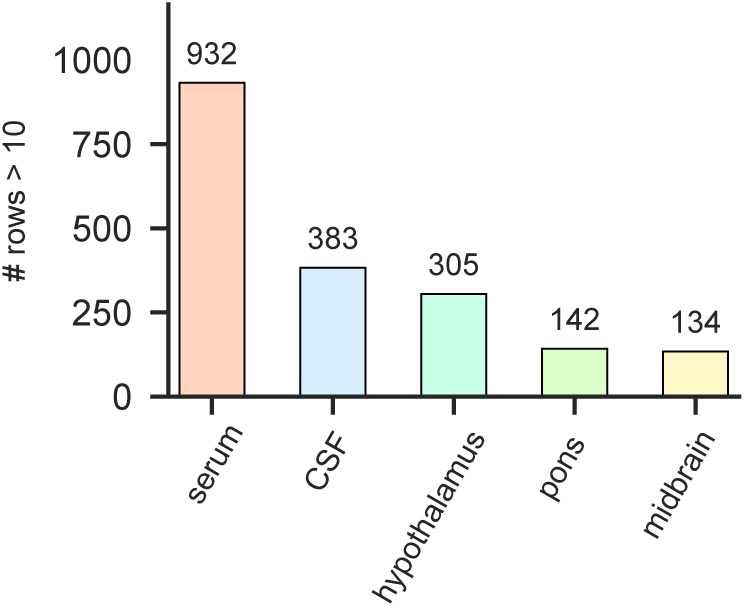
Brain-deposited autoreactive repertoire is restricted relative to serum and CSF. IgG was eluted from fresh brain sections and profiled by PhIP-seq to assess whether tissue reactivity reflected antigen-specific binding within brain rather than passive diffusion from blood or CSF. Bar plot shows the number of autoreactive peptides detected in each compartment at a fold-change enrichment cutoff >10. Serum contained 932 enriched peptides and terminal CSF (time point 2) contained 383 enriched peptides, whereas fewer enriched peptides were detected in brain regions (hypothalamus, 305; pons, 142; midbrain, 134), consistent with a markedly restricted brain-deposited autoreactive repertoire compared to serum and CSF.

**Extended Data Fig. 9.**
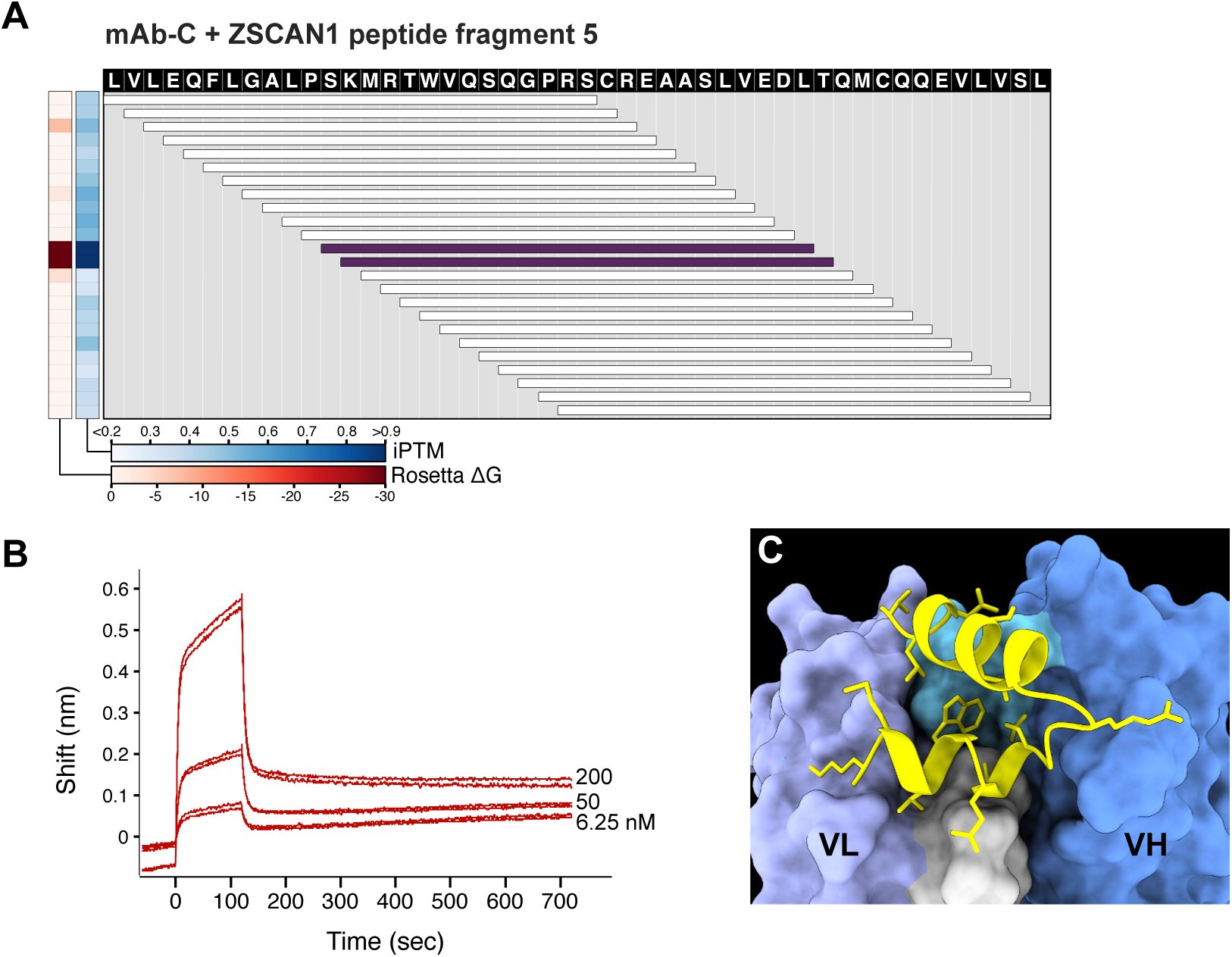
mAb-C binds to ZSCAN peptide fragment 5. **A)** Validation of PAIRIS binding predictions by biolayer interferometry (BLI) for mAb-C against ZSCAN1 peptide fragment 5. Heatmap displays iPTM scores (average interface confidence between peptide and both antibody chains) and Rosetta ΔG values for 25-mer peptides spanning ZSCAN1 peptide fragment 5 (aa 121-169, UniProt: Q8NBB4). Horizontal bars represent overlapping peptides spanning the indicated amino acid positions, with the color representing the sum of iPTM and Rosetta values. **B)** BLI sensorgram using displayed 48AA peptide and synthesized mAb-C. Reliable affinity estimation was limited by peptide insolubility. **C)** AlphaFold3-predicted structure shows the highest-scoring peptide-antibody complex, with sidechain atoms within 4Å displayed.

**Extended Data Fig. 10.**
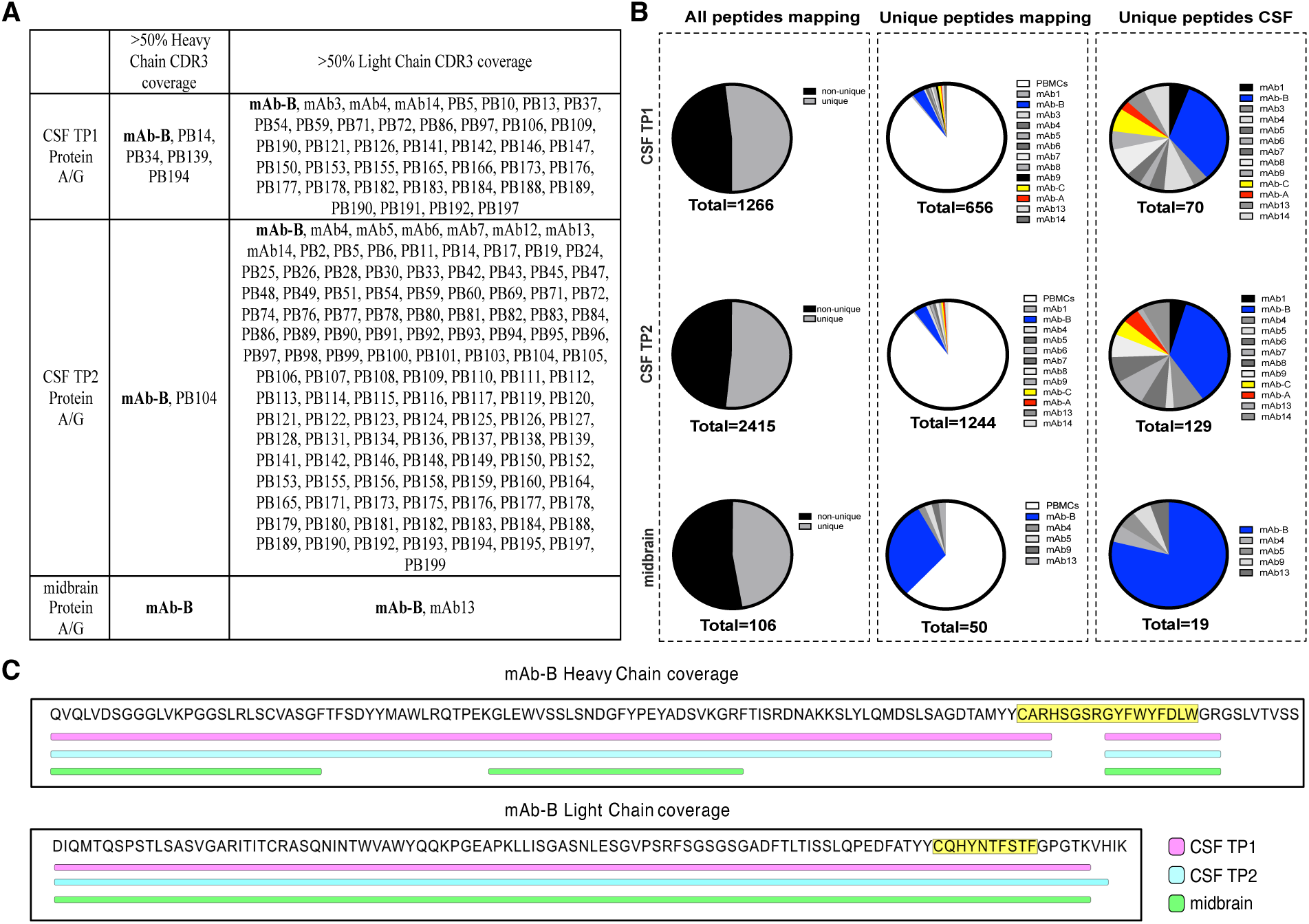
Ig mass spectrometry and peptide coverage of Clone B BCR. **A)** All peptides from protein A/G immunoprecipitation were mapped to the patient derived Ig database from either CSF or PBMCs. If at least 50% of the CDR3 region of the heavy or light chain were covered by a trypsin peptide from mass spectrometry, the antibody name is listed. Only mAb-B had >50% coverage in heavy and light chains in all samples. **B)** Unique peptides from trypsin digested CSF or midbrain lysate Protein A/G immunoprecipitates were mapped to the patient derived Ig database. Peptides that mapped uniquely vs non-uniquely (as in the case of many framework antibody regions) are shown on the first panel. Of the uniquely mapping peptides, the abundance of PBMC versus CSF derived antibodies are shown in the second panel. The last panel shows the uniquely mapping peptides aligning to the CSF only Ig database. **C)** Unique peptides identified by Protein A/G mass spectrometry from CSF or midbrain tissue lysate were aligned to a database of all patient derived CSF and PBMC BCRs identified from 10x sequencing. Clone B BCR showed the highest overall sequence level coverage, where bars represent at least one unique peptide mapping to the corresponding sequence. Yellow highlighted sequences indicate the CDR3 regions.

**Extended Data Fig. 11.**
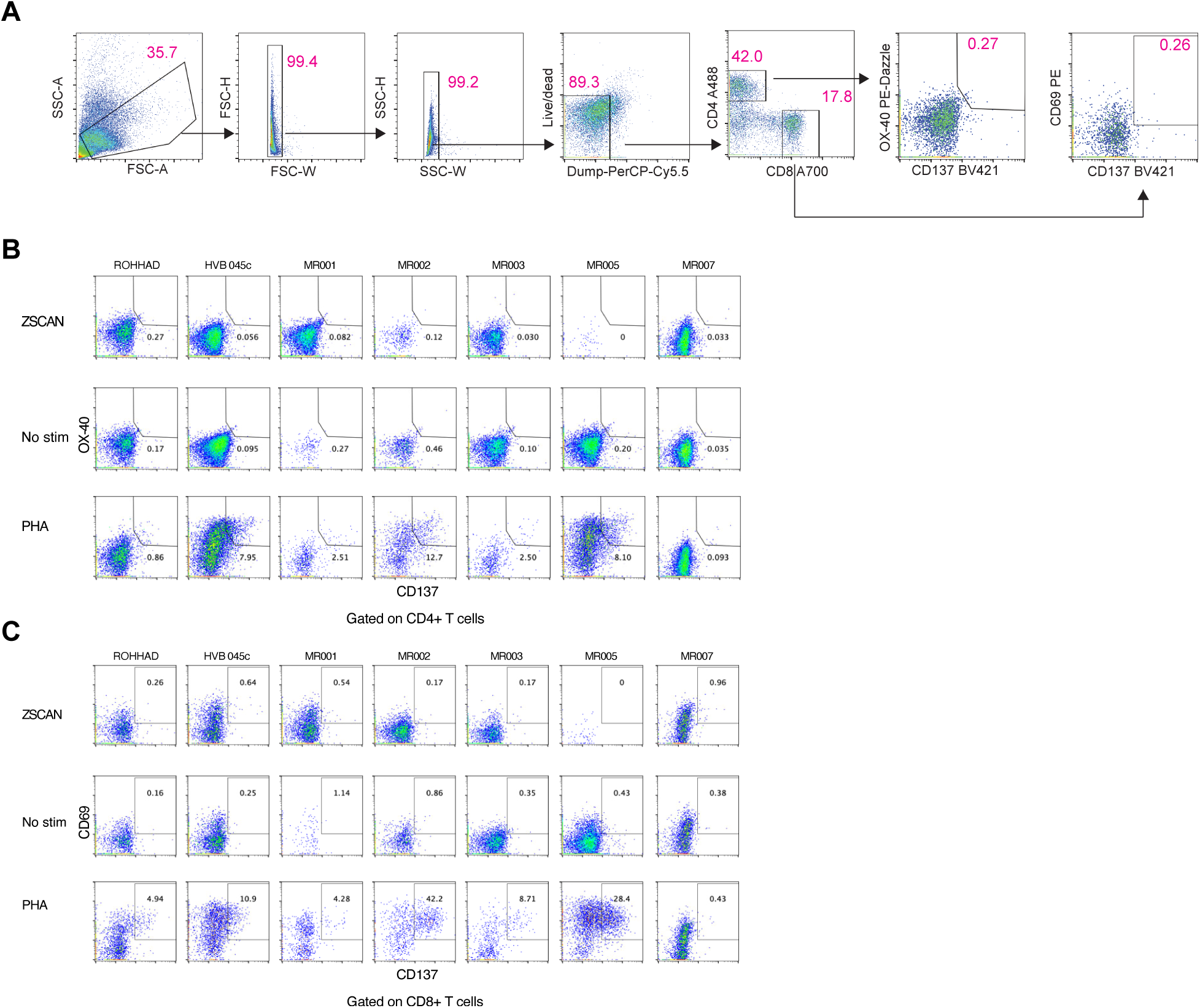
AIM Assay gating strategy and ZSCAN1-induced responses. **A)** Flow cytometry gating strategy for the activation-induced marker (AIM) assay. PBMCs were gated sequentially on lymphocytes (FSC-A vs SSC-A), singlets (FSC-H vs FSC-A and SSC-W vs SSC-A), live cells (Live/Dead eFluor 506–), and dump-negative cells (CD14/CD16/CD19–; PerCP-Cy5), followed by CD3+ T cells and separation into CD4+ and CD8+ subsets (CD4 vs CD8). AIM+ cells were then identified as CD137 (4-1BB)+OX40 (CD134)+ within CD4+ T cells and as CD137 (4-1BB)+CD69+ within CD8+ T cells (gates shown). **B)** Representative CD4+ AIM readouts (CD137 vs OX40) under no stimulation (0.2% DMSO), ZSCAN1 peptide pool stimulation, or PHA-L positive control, for the indicated samples/donors. **C)** Representative CD8+ AIM readouts (CD137 vs CD69) under the same stimulation conditions and samples/donors as in (B).

**Extended Data Fig. 12.**
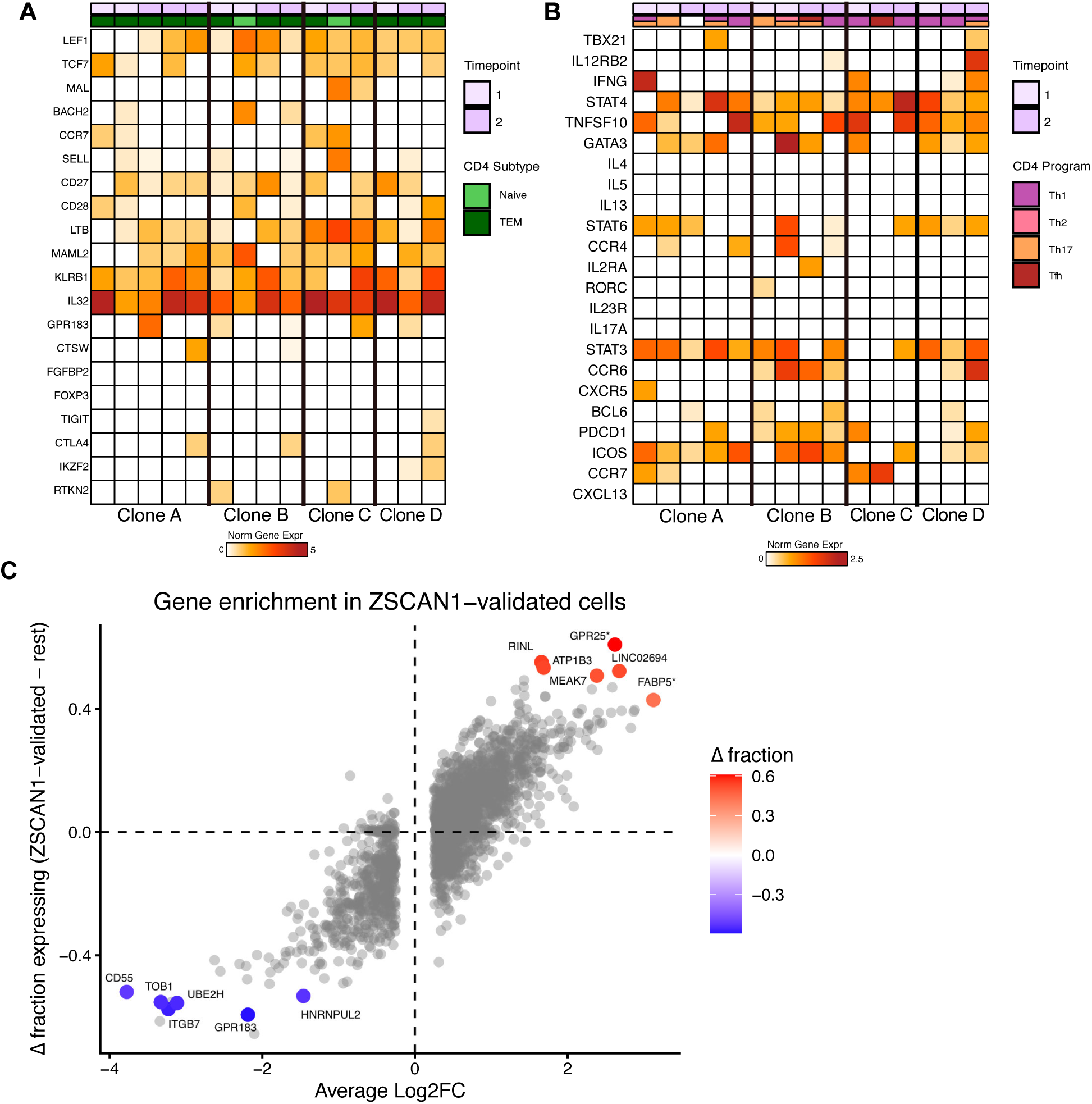
**CD4 T cell gene expression profiling**AUCell-based prediction of CD4 T-cell states for clones with ZSCAN1 autoreactivity, with heatmaps showing normalized expression of marker gene sets across timepoints for **A**) CD4 differentiation subtypes (Naive, TCM, TEM, Treg) and **B**) CD4 helper T-cell programs. **C**) Scatter plot showing the average log2 fold change versus the difference in fraction of cells expressing each gene in ZSCAN1-validated cells minus all other cells. Only genes expressed in >0.5 of either group are shown. Starred genes are significant at an adjusted p-value < 0.05. Genes in the top right are upregulated in ZSCAN1-reactive cells, while genes in the bottom left are absent or downregulated in ZSCAN1-validated cells.

